# Glymphatic function restored by *α*1-noradrenergic antagonism alleviates headache allodynia in mice

**DOI:** 10.64898/2026.04.24.720660

**Authors:** Adriana Della Pietra, Adisa Kuburas, Mathew Sevao, Tristen M. Castillo, Quinn K. Hanigan, Thomas L. Duong, Harold C. Flinn, Emilie H. Partridge, Murray A. Raskind, Jeffrey J. Iliff, Andrew F. Russo

## Abstract

Mild traumatic brain injury (mTBI) often leads to migraine-like post-traumatic headache (PTH), yet effective treatments are limited. Clinical and preclinical studies have shown that mTBI disrupts glymphatic transport of cerebrospinal fluid in the brain. We hypothesized that altered glymphatic transport might underlie facial allodynia commonly associated with migraine and PTH.

A closed-head impact model was used to induce mTBI in mice. Facial allodynia, a symptom of PTH and migraine, was evaluated using periorbital von Frey testing. Glymphatic influx was assessed using slice-based imaging of a fluorescent tracer injected into the cisterna magna.

Here we show that prazosin (PZN), an α1-noradrenergic receptor antagonist, restores glymphatic function and treats facial allodynia induced by calcitonin gene-related peptide (CGRP) and a nitric oxide donor in mice. In contrast, propranolol, a β-noradrenergic receptor antagonist, was ineffective. Even in the absence of mTBI, CGRP reduced glymphatic function and PZN was able to restore glymphatic function in the dorsal cortex. Importantly, the role of glymphatic function was confirmed by the lack of PZN efficacy in aquaporin-4 knockout mice.

These findings indicate that targeting α1-noradrenergic receptors to enhance glymphatic transport may offer a therapeutic strategy for treating migraine and PTH.

## Introduction

The global prevalence of new traumatic brain injuries was estimated at over 37 million per year in 2021^1^. TBI ranges in severity, from mild to severe, with mild traumatic brain injury (mTBI) accounting for 70-90% of cases^2^. mTBI is responsible for most of the TBI-related healthcare costs. Many mTBI patients develop post-traumatic headache (PTH), which often presents with migraine-like symptoms, including light/touch/sound sensitivity, nausea, vomiting, and impaired cognitive and psychosocial functions^3^. Studies vary on which phenotype is more prevalent^4,5^. Notably, headache persists in 41% of patients up to a year post-injury^6^. Several rodent models to study migraine and PTH have been developed. In this study, for PTH, we have used a closed head impact model of mTBI followed by injection of ordinarily subthreshold doses of the neuropeptide calcitonin gene-related peptide (CGRP) or the nitric oxide donor sodium nitroprusside (SNP). Both CGRP and SNP are well-documented triggers of migraine in people^7,8^ and drugs that target CGRP are approved for treating and preventing migraine (for review^9^).

We hypothesized that glymphatic dysfunction plays a critical role in PTH and migraine. The glymphatic system, introduced by Nedergaard and colleagues in 2012^10^, is a brain-wide network of perivascular pathways formed by astroglial cells. This system is responsible for the rapid exchange and clearance of cerebrospinal and interstitial fluid from the brain. Proper glymphatic function in rodent models is essential for clearing reactive oxygen species that are linked to neuroinflammation associated with migraine^11,12^, and PTH after TBI^3,13^. In particular, glymphatic function after TBI is reduced by ∼60% and remains impaired for at least a month^14^. Additional studies showed impaired clearance by glymphatic transport after closed-head impact TBI in rodents^15–17^. Studies in humans report MRI features proposed to reflect glymphatic dysfunction following TBI^18,19^, and whose presence predict the persistence of post-concussive symptoms including PTH^20^.

Glymphatic function requires astroglial aquaporin-4 (AQP4) water channels. The role of AQP4 channels has been demonstrated by genetic deletion of the AQP4 channel^10,14,21,22^, loss of perivascular AQP4 localization with deletion of *Snta1*^23^, and the pharmacological inhibition of AQP4^24^, which have all been shown to impair perivascular exchange. However, to our knowledge no studies have looked at mTBI-induced behavor in mouse AQP4 deficient mouse models.

Importantly, glymphatic dysfunction may elevate the levels of CGRP in the cerebrospinal fluid (CSF). CGRP is a multifunctional neuropeptide that is most notable as a key player in migraine pathogenesis^25,26^. A recent paper reported that solutes, including CGRP, can be directly transported from the cortex to the trigeminal ganglion via the CSF, forming a non-synaptic communication pathway between the central and peripheral nervous systems^27^. This transport to the trigeminal ganglion was increased following experimentally induced cortical spreading depolarization, which is associated with the aura phase of migraine and commonly occurs after TBI^28–30^. Interestingly, AQP4 knockout (KO) mice, which exhibit reduced glymphatic function, showed an even greater impairment following closed-head impact TBI^14^. This observation is consistent with decreased AQP4 expression, and astrocytic end-foot depolarization reported by Ren et al. starting three days post-injury^31^. We propose that reduced glymphatic clearance post-mTBI leads to the accumulation of solutes such as CGRP that can cause migraine or migraine-like PTH.

Tackling PTH from a different perspective, sleep disruption is also a known migraine trigger and is frequently reported following mTBI, whilst sleep is a commonly reported migraine abortive^32^. Sleep-wake differences in glymphatic function are regulated by central noradrenergic tone^33^. It was previously established that local infusion in the cisterna magna of a cocktail of noradrenergic antagonists consisting of the α1 antagonist prazosin (PZN), the α2 receptor antagonist atipamezole, and the β antagonist propranolol (PPL) increased perivascular CSF influx and interstitial solute clearance in awake animals to levels observed during sleep or anesthesia^34^. For the present study, we evaluated the effects of PZN and PPL. We selected PZN because it is used to treat trauma nightmares in the setting of post-traumatic stress disorder (PTSD)^35^ and importantly, a small preliminary clinical trial (PZN n=30, placebo n=16) reported that PZN reduced headache frequency in Veterans diagnosed with PTH^36^. PZN was also an attractive candidate since it is a relatively safe drug that has been used by millions of people in the decades since it was initially developed to treat hypertension in the 1970s^37^. PPL is currently used as a preventive drug for migraine^38^.

In this study, we demonstrate that enhancing glymphatic transport by treating mice with the α1-noradrenergic receptor antagonist PZN, but not the b-noradrenergic antagonist PPL, can rescue facial allodynia caused by two different migraine triggers in mice across both PTH and primary migraine mouse models. We establish a causal relationship between glymphatic system function and the alleviation of allodynia by demonstrating that PZN efficacy requires AQP4 water channels. These results indicate that modulation of the glymphatic system may prove beneficial for treating migraine and PTH in patients.

## Methods

### Animals

An equal number of male and female WT outbred CD1 mice were obtained from Charles River Laboratories (Roanoke, IL) and WT C57BL/6J mice from Jackson Laboratories at 9 weeks of age. Mice were housed under 12-h dark/light cycles, with 4-5 mice per cage, ad libitum access to food and water, and ambient temperature of 22°C. Mice were tested after at least one week of acclimation in the animal facility. All experiments were from at least two independent cohorts, unless specified differently in the figure legends. The parental *Aqp4^+/-^*transgenic KO mice (B6(Cg)-Aqp4<tm1.1Tsna>) were obtained originally from RIKEN BRC, Japan. The mice were transferred from the Iliff lab to Iowa with permission from RIKEN. Heterozygous mice were crossed among themselves to generate homozygous *Aqp4^−/−^* KO and nontransgenic littermates. The *Aqp4^−/−^* mice were maintained as a homozygous colony and the nontransgenic mice were bred with C57BL/6J mice and maintained as WT controls in parallel breeding cages. For behavior studies, mice were selected between 10 to 12 weeks of age. All experiments were performed between 8 AM and 5 PM with randomized mice and investigators blinded to injury and drug treatments. AK was the sole investigator aware of group allocation throughout the experiments. All other authors conducted the experiments, assessed outcomes, and analyzed the data while blinded. Group allocation was revealed only upon completion of each experimental group. All animal procedures performed in this study were approved by the University of Iowa Animal Care and Use Committee and follow the rules set by the National Institutes of Health, policies of the International Association for the Study of Pain and ARRIVE guidelines.

### Drug administration

PZN (Sigma-Aldrich, St Louis, MO) was administered in the drinking water at 45 mg L^−1^, which is estimated to be equivalent to 7.5 mg kg^−1^ day^−1^ for each mouse. PPL (Sigma-Aldrich, St Louis, MO) was administered in the drinking water at 240 mg L^−1^ day^−1^ for chronic administration experiments, which is estimated to be equivalent to 40 mg kg^−1^ day^−1^ for each mouse. In acute administration experiments, PZN was prepared in Dulbecco PBS (Hyclone) and injected i.p. 0.3 mg kg^−1^ or 3 mg kg^−1^ and PPL at 1.7 mg kg^−1^ 1 h before behavioral testing. The initial acute doses for PZN and PPL were calculated to match the estimated amount consumed over 1 h in the chronic drinking-water paradigm, assuming constant water intake evenly distributed across the day. Thus, for PZN: 7.5 mg kg^−1^ day^−1^ / 24 h day^−1^= 3 mg kg^−1^ h^−1^; and for PP:, 40 mg kg^−1^ day^−1^ / 24 h day^−1^= 1.7 mg kg^−1^ h^−1^. The following compounds were prepared in Dulbecco PBS (Hyclone) and injected i.p.: 0.1 mg kg^−1^ or lower rat αCGRP (Sigma-Aldrich, St Louis, MO), 0.25 mg kg^−1^ SNP (Sigma) in the same volume as vehicle. All injections were performed by AK. All injections were i.p. with a 0.3 x 13 mm needle (insulin needle).

### Closed head impact repetitive mTBI

Closed head impact mTBI was performed with one injury per day for 3 consecutive days following a previously described protocol^39^. In brief, mice were anesthetized with 5% isoflurane for 2 min, then placed on a double-layer foam cushion to prevent head rotation. The setup consists of a vertically positioned fiberglass tube, maintained just against the surface of the mouse head. A stainless-steel weight was released from the top of the tube to strike the upper-right hemisphere of the head, targeting the temporal area. For CD1 mice, the tube is 80 cm long with an inner diameter of 13 mm and the metal weight is 30 g. For the smaller C57BL/6J mice, the tube length is 50 cm with the same 13 mm diameter and a metal weight of 21 g. After the impact, the mice were placed back in their cages and monitored until awake (within a few min). Sham mice were anesthetized but not subject to weight-drop. Both mice groups gained weight at the same rate and a previous study has confirmed that there is no detectable skull fracture or brain hemorrhage after this trauma^39^.

### Intracisternal tracer infusion for assessment of static glymphatic flow

Intracisternal infusion of a tracer to track the glymphatic flow was performed as previously described^10^. A 10 μl Hamilton syringe (Hamilton 80001) connected to a 30-gauge needle via PE-10 tubing was prepared by filling the line with saline and backfilling with tracer solution of Texas Red 0.5% dissolved in artificial CSF immediately before injection. Mice were anesthetized with i.p. administration of ketamine/xylazine (87.5 mg kg^−1^/12.5 mg) for deep anesthesia and injected with i.p. CGRP (0.1 mg kg^−1^) or vehicle. Animals were then positioned in a stereotaxic frame, and the skin on the posterior neck was cleared of hair using an electric razor. A midline incision was made to expose the underlying muscle, which was carefully retracted to reveal the atlanto-occipital membrane over the cisterna magna. The needle was inserted into the cisterna magna such that the bevel passed through the membrane, and VetBond (Thermo Fisher Scientific, MA, USA) was applied to stabilize the needle in place. Tracer solution was infused using a syringe pump at a rate of 1 μl /min for a total of 10 min, delivering 10 μl in total The interval between i.p. CGRP or vehicle administration and tracer infusion corresponded to the time required to prepare the animal for infusion as described above. There was no significant difference between groups in the time between i.p. CGRP (21.8 ± 1.4 min) or vehicle (20.1 ± 1.1 min) administration and tracer infusion (mean ± SEM). Injection success was verified by post-mortem visualization of tracer along perivascular pathways. Mice remained under anesthesia throughout the ∼50 min procedure, with additional doses given as needed to a maximum of 4 injections per mouse determined by the pedal withdrawal reflex. Body temperature was maintained between 36–38 °C using a heating pad and heat lamp. At the end of the experiment, mice were perfused transcardially with 4% paraformaldehyde in PBS. Whole brains were post-fixed overnight in 4% paraformaldehyde, cryoprotected in 30% sucrose until they sank to the bottom of the container, embedded in OCT, and stored at –80 °C until sectioning. Mice were excluded from the study if they presented with fight wounds, severe dermatitis, or blood around the ears from excessive grooming. Additionally, mice that died during tracer infusion, or whose imaged brains showed no fluorescence, indicating a failed injection, were removed from statistical analysis.

All image analysis was performed by Fiji (ImageJ) for quantifying the mean grey values of different areas in brain slices. For each mouse 3 slices in the rostral and caudal brain regions were selected and analyzed as repeated measures (sections 1 at bregma + 0.25, 3 at bregma - 1.25, and 5 at bregma - 2.75, as previously described^40^). The hippocampus was present only in sections 3 and 5.

### Periorbital von Frey

Periorbital allodynia was evaluated by the up and down protocol as previously published^39^. To summarize, CD1 mice were acclimated for 1 week and C57BL/6J mice for 2 weeks to their own polycoated paper cup (4 oz. paper cups; 6.5 cm top diameter, 4.5 cm bottom diameter, 72.5 cm length) positioned on top of a plastic holder. During the acclimation, mice were habituated to filament D (0.07 g). Mice were considered acclimated when filament D could touch the periorbital area without a reaction from the mouse.

Each testing started with filament D. If the mouse did not react, the heavier filament E was applied, proceeding until filament H (1 g). Once the mouse responded, the lighter filament (e.g. C after D) was used, going down until filament A (0.008 g). A pattern was recorded for each response or non-response. The same approach was used with up to 5 filaments after the first change in pattern. The last filament reached in the pattern and the pattern itself were used to calculate the 50% threshold, according to an established calculation^41^. Mice were excluded from testing either for having fight wounds, severe dermatitis or blood on bilateral ear bases provoked by excessive grooming. Mice exhibiting excessive activity that prevented reliable assessment (or testing) on specific test days were excluded from analysis for those time points only.

### Experimental design

Before each behavioral assay, mice were habituated to the testing room for 1 h. A baseline was measured for each mouse the day before mTBI. The mice were then divided into groups to ensure comparable baseline measurements. All the mice in the same cage had repetitive mTBI or sham and all shared the same drinking water with or without drug. Drugs in the drinking water were administered on the first day of mTBI immediately following the first injury or 2 weeks post-mTBI. The drug was removed after 6 weeks for 2-4 weeks then reintroduced for 2 more weeks. Test time-points were every 2 weeks after mTBI. Mice were tested 30 min after CGRP or SNP i.p. injections for periorbital sensitivity. For glymphatic measurements, CGRP was injected 30 min before intracisternal injection of Texas Red-conjugated dextran. In the experiment where PZN was administered on day 1 of mTBI in CD1 mice, one cohort was tested on day 49 and the other two on day 42 due to internal lab scheduling. These data were combined in the graph because the cohorts exhibited identical behavior. All other measurements were taken at the indicated days.

### Statistical analysis

Power analysis was performed for group sample size calculations by ClinCalc.com. The estimated effect size for this study was a 50% reduction in periorbital tactile hypersensitivity, based on data from similar previous studies^39^. With an alpha of 0.05 and a power of 0.80, the projected sample sizes required were 12-14 mice per group for tactile allodynia. Based on a previous paper^10^, approximately 6 mice per group were used for the glymphatic fluorescence analysis.

Data were analyzed using GraphPad Prism version 10.6.1 for MacBook (GraphPad Software, Boston, Massachusetts USA). When powered for sex differences, data were also represented in male and female groups to show sex-dependent differences. All data are represented as mean ± 95% confidence interval. Histogram data were analyzed for normal distribution by Shapiro-Wilk test and Kolmogorov-Smirnov test. Data failing any of these tests were considered non-normally distributed and analyzed by non-parametric statistical tests. Normally distributed histograms were analyzed with one-way ANOVA with post-hoc Tukey’s multiple comparison test. Longitudinal data were analyzed with a two-way repeated measure ANOVA or mixed effects analysis with post-hoc Tukey’s multiple comparison test. All statistics are specifically reported in Supplementary Table 1. All data are available on request.

## Results

### Prazosin Concomitantly with mTBI rescues CGRP-induced periorbital sensitivity in CD1 mice

To test whether PZN might alleviate PTH symptoms, we used mTBI-induced periorbital allodynia elicited by subthreshold CGRP as a model with outbred CD1 mice. The experimental protocol for PZN administration is outlined in Figure 1A. Closed-head impact mTBI was performed once a day for three consecutive days. PZN was administered in the drinking water immediately after the first injury and maintained for 6 weeks. Mice were tested for periorbital sensitivity using von Frey filaments at baseline one day before the first mTBI. Senstivity during the acute phase was assessed one day after the last mTBI. In the von Frey assay, increased sensitivity is measured as decreased threshold. Chronic sensitivity to a normally subthreshold trigger was then tested at approximately 2 week intervals beginning 2 weeks after injury, with testing at 30 min after an intraperitoneal (i.p.) injection of subthreshold CGRP (subCGRP) (0.01 mg/kg) or vehicle. After 6 weeks, PZN was removed and the mice retested after 2 weeks without PZN, followed by testing again after reintroduction of PZN for 2 weeks.

**Figure 1.**
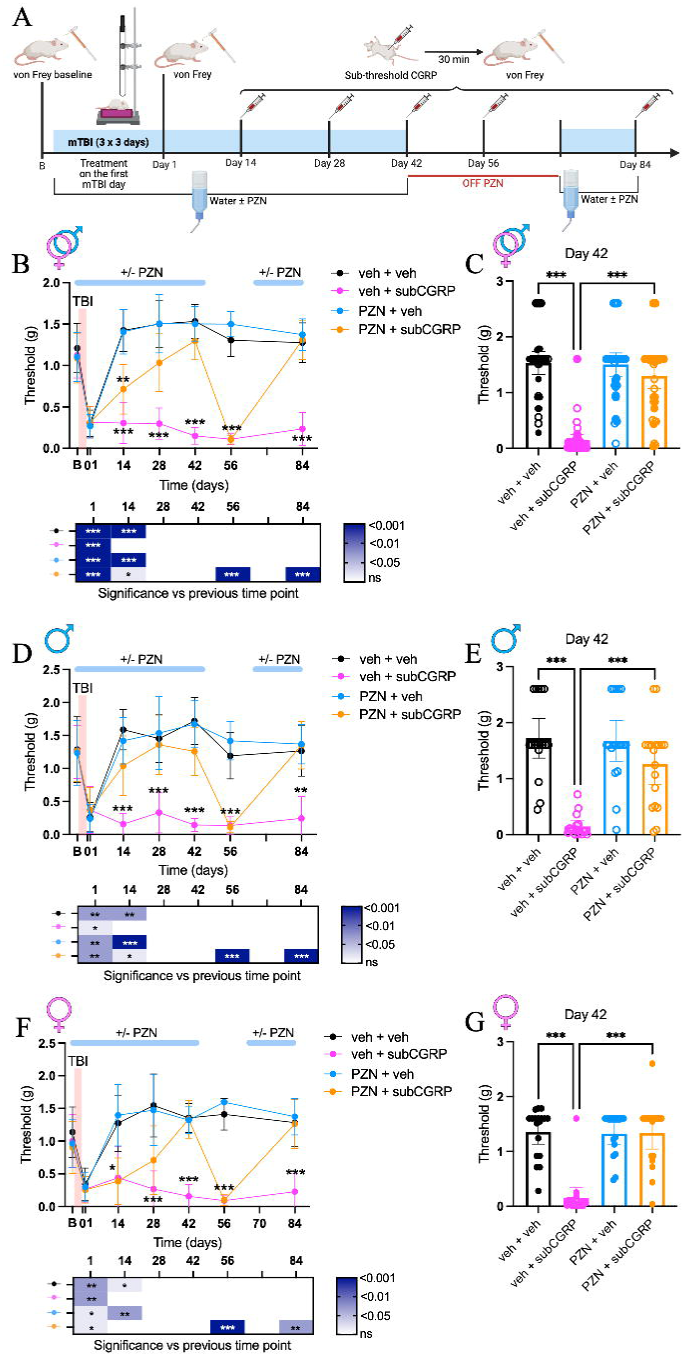
Treatment with PZN concomitantly with mTBI rescues subthreshold CGRP-induced periorbital allodynia in CD1 mice. **(A)** Protocol schematic. Periorbital allodynia was assessed at baseline (B) and at the indicated time points after mTBI without any treatment (day1), and then 30 min after i.p. administration of vehicle (veh), or subthreshold CGRP (0.01 mg kg^−1^). PZN treatment was started on the first day of mTBI in drinking water and continued for 6 weeks. PZN was then removed for 4 weeks and reintroduced for 2 weeks. **(B)** Longitudinal data for all mice with significance between each treatment compared to veh + veh are reported on the lines of the graph. The significance within each group for a time point compared to the previous time point is reported in the heatmap. **(C)** Scatter plot of individual mice from the longitudinal plot at day 42 post-mTBI. Open symbols for males and closed symbols for females. **(D), (E)** Longitudinal and day 42 scatter plot data from male mice of panels b and c. **(F), (G)** Longitudinal and day 42 scatter plot data from female mice of panels b and c. Data shown in all histograms at day 42 include an additional cohort tested at day 49. Since no differences were observed between the two time points, the data were pooled. Statistics in Supplementary Table 1.

At day 1 post-mTBI, all mice showed an acute increase in the periorbital sensitivity from baseline in the presence or absence of PZN. By 2 weeks (day 14), this acute sensitivity was resolved for all mice. During the chronic sensitivity phase, a subCGRP injection dose that was insufficient to induce allodynia in sham uninjured mice^39^, produced a marked increase in sensitivity in mice on water with vehicle. In contrast, mice on water with PZN exhibited a partial recovery to subCGRP by day 14. This recovery gradually improved to a complete rescue at day 42, when the sensitivity returned to the original baseline level (Figure 1B, C). We note that data were collected on day 49 for one cohort of mice and on day 42 for the other two cohorts. Since the day 42 and 49 data were statistically the same, the data were combined. When compared by sex, male mice were rescued by PZN faster than females. At day 14, males were fully rescued (Figure 1D, E), while females were not fully rescued until day 42 (Figure 1F, G). Thus by 6 weeks (day 42), PZN fully rescued subCGRP-induced periorbital allodynia in all mice post-mTBI.

To address whether the rescue by PZN was permanent or if continued treatment was required to prevent the allodynic response, we removed PZN from the drinking water after day 42 (Figure 1A). Two weeks after PZN removal, at day 56, periorbital sensitivity upon subCGRP administration was increased to the same level as the vehicle control mice that had not previously received PZN (Figure 1B). Upon reintroduction of PZN, the protective effect of PZN was fully restored by day 84, which was a faster recovery compared to the initial treatment. The same effect was observed in both male (Figure 1D) and female mice (Figure 1F). Thus, continuous PZN treatment is needed to rescue subCGRP-induced facial allodynia in CD1 mice after mTBI.

### Prazosin rescues CGRP-induced periorbital sensitivity in C57BL/6J mTBI mice

Since C57BL/6J is a standard strain used by many labs, we examined PZN effect on mTBI-induced periorbital allodynia using the same protocol as with CD1 mice (Figure 2A). All mice had increased acute periorbital sensitivity at day 1. As with CD1, subCGRP caused periorbital sensitivity in the absence of PZN and was rescued by PZN in the drinking water (Figure 2B, C). One difference from CD1 mice was that PZN fully rescued subCGRP sensitivity earlier by day 14. There was no sex-difference observed in these mice (Figure 2D-G). As with CD1, discontinuation of PZN led to increased periorbital sensitivity within 2 weeks and was fully restored after 2 weeks, regardless of sex (Figure 2B-G). To summarize, PZN treatment is effective when administered on the first day of mTBI in C57BL/6J mice similar to CD1 mice and continuous PZN treatment is needed to rescue subCGRP-induced periorbital allodynia.

**Figure 2.**
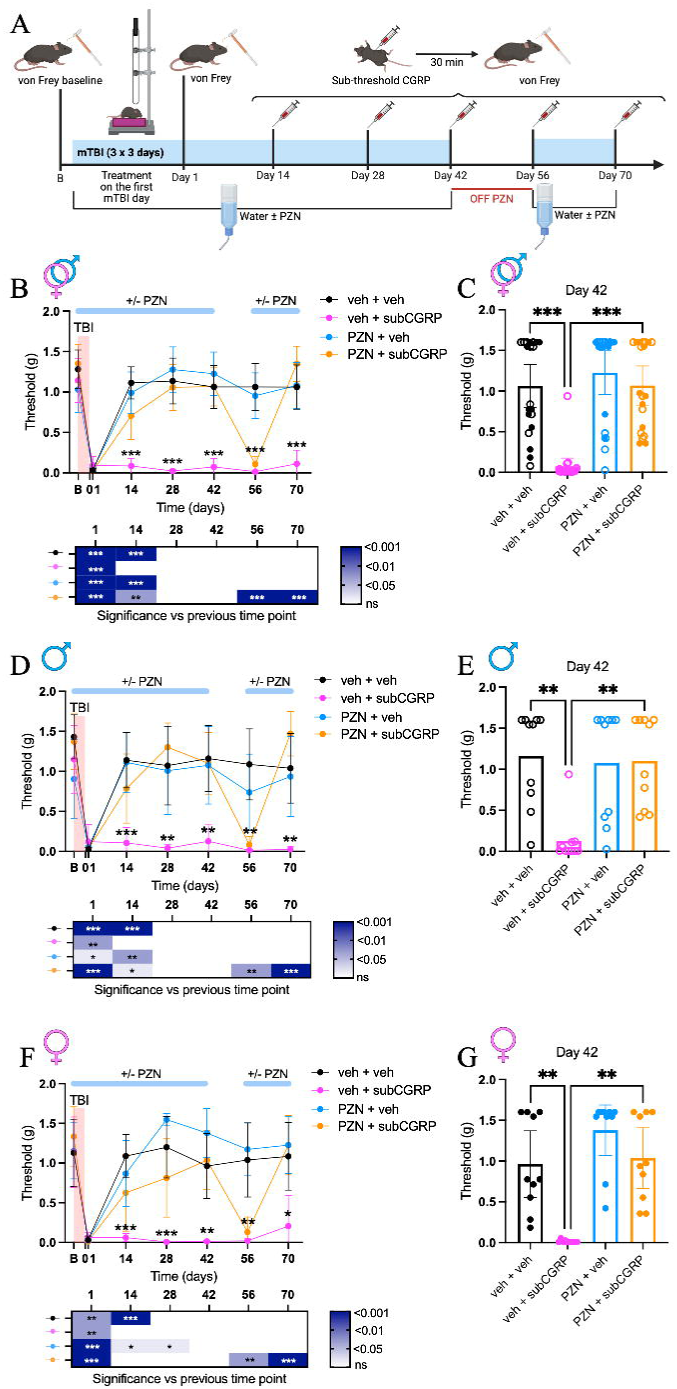
Treatment with PZN concomitantly with mTBI rescues subthreshold CGRP-induced periorbital allodynia in C57BL/6J mice. **(A)** Protocol schematic. Periorbital allodynia was assessed at baseline (B) and at the indicated time points after mTBI without any treatment (day1), and then 30 min after i.p. administration of vehicle (veh), or subthreshold CGRP (0.01 mg kg^−1^), as described in Figure 1. **(B)** Longitudinal data for all mice with significance between each treatment compared to veh + veh are reported on the lines of the graph. The significance within each group for a time point compared to the previous one is reported in the heatmap. **(C)** Scatter plot of individual mice from the longitudinal plot at day 42 post-mTBI. Open symbols for males and closed symbols for females. **(D), (E)** Longitudinal and day 42 scatter plot data from male mice of panels b and c. **(F), (G)** Longitudinal and day 42 scatter plot data from female mice of panels b and c. Statistics in Supplementary Table 1.

### Acute prazosin treatment does not rescue CGRP-induced periorbital allodynia in mTBI mice

We then asked whether acute PZN treatment might be sufficient for rescuing allodynia. The mice were tested 5-7 weeks post-mTBI at a point when we knew that chronic PZN treatment had been effective in parallel cohorts. PZN was injected i.p. 30 min before the subCGRP trigger then tested for periorbital sensitivity 30 min after subCGRP (1h post-PZN) (Figure 3A). A systemic dose of PZN of 0.3 mg kg^−1^ i.p. was estimated to be comparable to the amount ingested over 1 h in the drinking water (see Methods). This dose did not cause any apparent discomfort to the mice based on facial features (Figure 3B), but did not rescue subCGRP sensitization (Figure 3C). However, PZN on its own appeared to increase sensitivity, although this did not reach statistical significance. The lack of rescue was seen in both male and female mice (Figure 3D, E). A higher dose of 3 mg kg^−1^ i.p. PZN was also tested, although this dose induced apparent pain in mice based on squinting 1 h after administration (Figure 3F). At this dose, all mice given PZN had increased periorbital sensitivity, as did mice given subCGRP without PZN (Figure 3G-I). Thus, acute systemic administration of PZN 30 min before subthreshold CGRP was not sufficient to treat the periorbital allodynia in mTBI mice.

**Figure 3.**
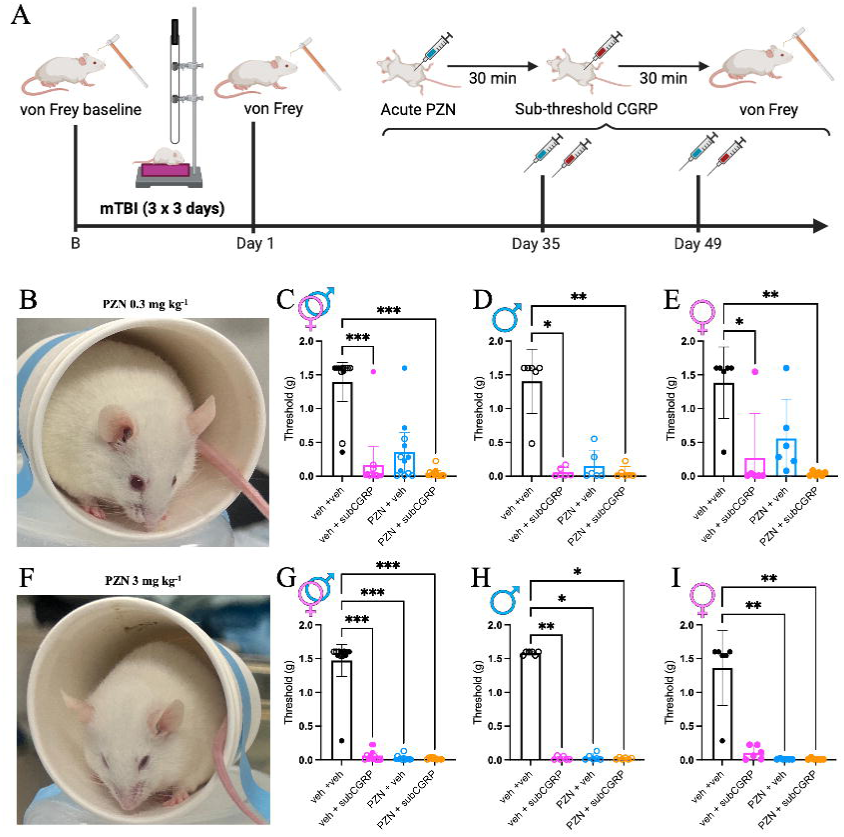
Acute PZN treatment does not rescue subthreshold CGRP-induced periorbital allodynia after mTBI. **(A)** Protocol schematic. Periorbital allodynia was assessed at baseline (B) and at the indicated time points after mTBI without any treatment (day1). Then, 1h after after i.p. administration of vehicle (veh), or acute PZN (3 mg kg^−1^ or 0.3 mg kg^−1^) and 30 min after i.p. administration of vehicle, or subthreshold CGRP (0.01 mg kg^−1^). **(B)** Picture of a mouse 1 h post i.p. injection with PZN 0.3 mg kg^−1^. **(C)** Scatter plot of all mTBI mice tested 1 h post-PZN (0.3 mg kg^−1^) or veh and 30 min post subCGRP (0.01 mg kg^−1^) or veh. Open symbols for males and closed symbols for females. **(D)** Scatter plot of male mTBI mice from panel c. € Scatter plot of female mTBI mice from panel c. **(F)** Picture of a mouse 1 h post i.p. injection with PZN (3 mg kg^−1^). The mouse shows non-evoked pain with squinting behavior. **(G)** Scatter plot of all mTBI mice tested 1 h post-PZN (3 mg kg^−1^) or veh and 30 min post subCGRP (0.01 mg kg^−1^) or veh. Open symbols for males and closed symbols for females. **(H)** Scatter plot of male mTBI mice from panel g. **(I)** Scatter plot of female mTBI mice from panel g. Statistics in Supplementary Table 1.

### PPL does not rescue CGRP-induced periorbital allodynia after mTBI

In addition to PZN, the noradrenergic antagonist cocktail used to enhance glymphatic function also contained a β-noradrenergic receptor antagonist, PPL^34^. Interestingly, PPL is commonly used as a migraine prophylactic drug^38^. To test whether PPL might also prevent subCGRP-induced allodynia following mTBI, mice were given PPL in their drinking water immediately after the first impact as done with the PZN protocol (Figure 4A). Mice were tested with periorbital von Frey at baseline and at the acute phase (day 1) post-mTBI. All mice showed an acute periorbital sensitivity response compared to baseline. Mice were tested for subCGRP sensitivity at day 14. In contrast to the rescue observed with PZN administration, mice on PPL did not show any recovery in sensitivity by day 14 (Figure 4B, C). Even worse, the mice drinking PPL showed a significant increase in periorbital sensitivity compared to vehicle, indicating a possible pro-allodynic effect of chronic PPL. The basis of this apparent effect is not known but appears to be rather variable (Figure 4C). For ethical reasons, the experiment was stopped after this first time point. Thus, chronic PPL did not rescue subCGRP-induced periorbital allodynia in mTBI mice by day 14 and in a subset of mice PPL appears to induce periorbital allodynia.

**Figure 4.**
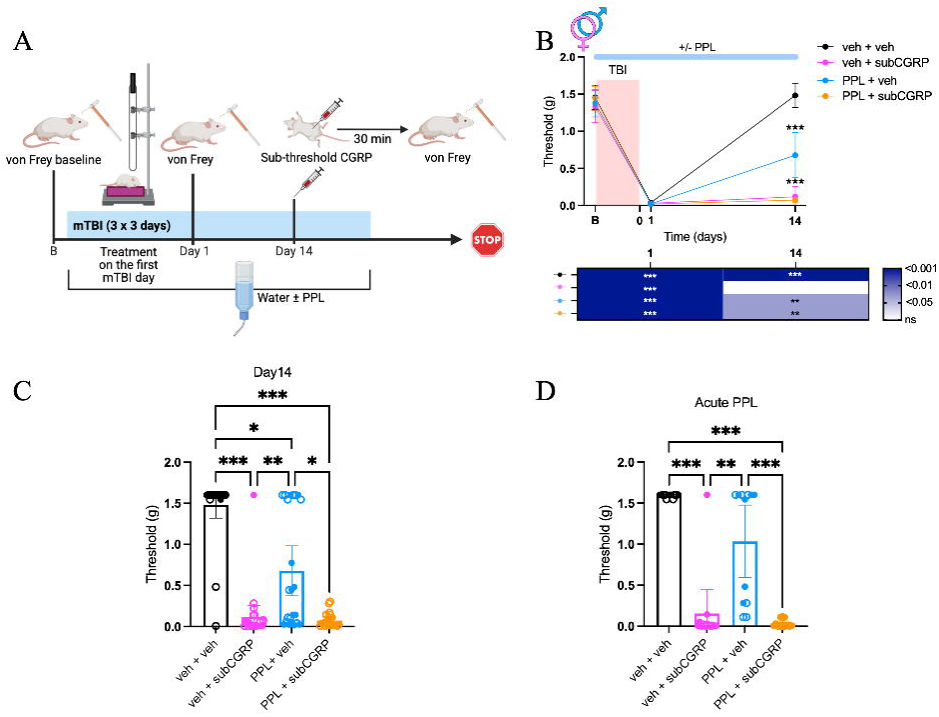
Acute or chronic PPL treatment does not rescue subthreshold CGRP periorbital allodynia in mTBI mice. **(A)** Protocol schematic. Periorbital allodynia was assessed at baseline (B) and at the indicated time points after mTBI without any treatment (day1), and then 30 min after i.p. administration of vehicle (veh), or subthreshold CGRP (0.01 mg kg^−1^). PPL treatment was started on the first day of mTBI in drinking water. After the day 14, the experiment was stopped for ethical concerns. **(B)** Longitudinal data for all mice with significance between each treatment compared to veh + veh are reported on the lines of the graph. The significance within each group for a time point compared to the previous one is reported in the heatmap. **(C)** Scatter plot of individual mice at day 14 post-mTBI from the longitudinal plot. Open symbols for males and closed symbols for females. **(D)** Scatter plot of individual mice tested 1 h post-acute PPL (1.7 mg kg^−1^) or veh and 30 min post subCGRP (0.01 mg kg^−1^) or veh. A separate cohort of mice was used for the acute experiment with testing at 63 days following mTBI. Open symbols for males and closed symbols for females. Statistics in Supplementary Table 1.

To match the PZN experiments, we also tested whether acute PPL treatment might rescue subCGRP-induced periorbital allodynia after mTBI. In a separate cohort of mice from the chronic PPL study, we administered a systemic dose of PPL (1.7 mg kg^−1^) i.p. that was estimated to be comparable to the amount ingested over 1 h in the chronic drinking-water studies (see Methods). This dose did not significantly increase periorbital sensitivity on its own but also did not rescue subCGRP allodynia (Figure 4D). Overall, these findings indicate that the rescue of subCGRP-induced allodynia in mTBI mice is mediated via α1 and not b-noradrenergic receptors.

### Prazosin administered after mTBI rescues CGRP-induced periorbital sensitivity

Given that most patients with mTBI will be treated after and not before mTBI, we next evaluated the therapeutic potential of PZN when administered following the last of multiple mTBIs. The protocol was similar to the earlier concomitant treatment paradigm with CD1 mice (Figure 1A), except PZN was given 2 weeks after the last day of mTBI and a sham, uninjured group was included (Figure 5A). At 2 weeks post-mTBI, sham mice received i.p. injections of either subthreshold CGRP (0.01 mg kg^−1^) or full dose CGRP (0.1 mg kg^−1^) 30 min before testing, while mTBI mice received vehicle or subthreshold CGRP. All mice were given PZN in their drinking water for 6 weeks, followed by a 2-week PZN withdrawal, and then reintroduced to PZN for an additional 2 weeks. Periorbital allodynia thresholds were assessed every 2 weeks.

**Figure 5.**
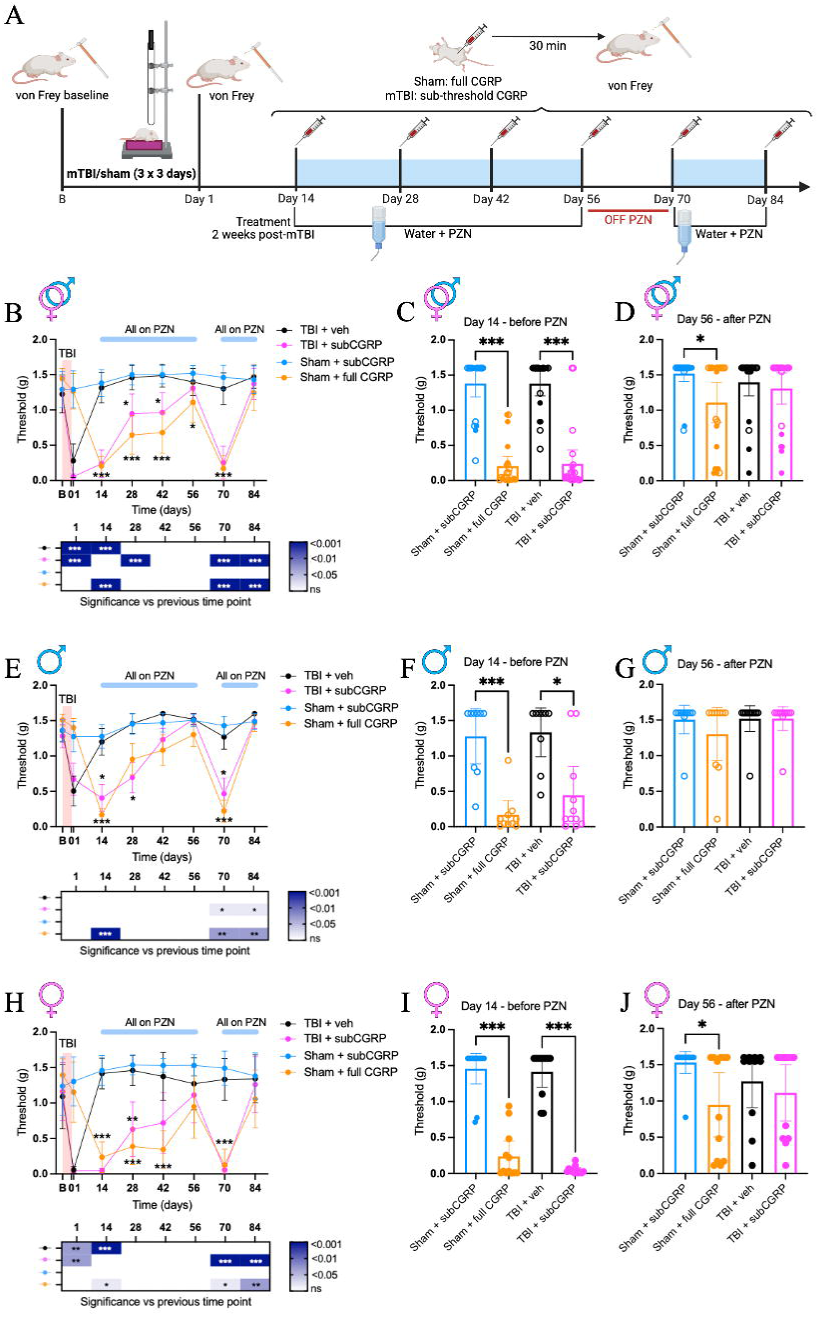
PZN treatment post-mTBI rescues subthreshold and full-dose CGRP periorbital allodynia. **(A)** Protocol schematic. Periorbital allodynia was assessed at baseline (B) and at the indicated time points after mTBI/sham without any treatment (day1), and then 30 min after i.p. administration of vehicle (veh), or subthreshold CGRP (subCGRP, 0.01 mg kg^−1^) or full dose CGRP (full CGRP, 0.1 mg kg^−1^). PZN treatment was started at 2 weeks post-mTBI in drinking water of all mice and continued for 6 weeks up to day 56. PZN was then removed for 2 weeks and reintroduced for 2 more weeks. **(B)** Longitudinal data for all mice with significance between treatments: sham + subCGRP vs sham + full CGRP, and TBI + veh vs TBI + subCGRP. The significance within each group for a time point compared to the previous one is reported in the heatmap. **(C)** Scatter plot of individual mice at day 14 post-mTBI, before PZN administration, from the longitudinal plot. Open symbols for males and closed symbols for females. **(D)** Scatter plot of individual mice at day 56 post-mTBI, after PZN administration, from the longitudinal plot. Open symbols for males and closed symbols for females. **(E)-(G)** Longitudinal and day 14 and day 56 scatter plot data from male mice of panels b-d. **(H)-(J)** Longitudinal and day 14 and day 56 scatter plot data from female mice of panels b-d. Statistics in Supplementary Table 1.

As expected, mTBI mice had a significant increase in their periorbital sensitivity from baseline to day 1 post-mTBI during the acute phase (Figure 5B). By day 14 post-mTBI, the acute effect had worn off. At this time point, before PZN administration, sham mice given subthreshold CGRP and mTBI mice given vehicle showed von Frey sensitivities similar to baseline, while sham mice given full dose CGRP and mTBI mice given subCGRP displayed increased sensitivity, as expected (Figure 5B, C). PZN was then administered in drinking water to both sham and mTBI mice for 6 weeks up to day 56. mTBI mice injected with subCGRP showed partial recovery by day 28 and full recovery by day 56 (Figure 5B, D). So, 6 weeks of continuous PZN treatment fully rescued subCGRP-induced periorbital sensitivity even when treatment was initiated post-mTBI.

An unexpected observation was that PZN also reduced the allodynia in sham mice that had not experienced mTBI, but had been treated with a full dose of CGRP (Figure 5B, D). At day 56, the allodynia was clearly reduced, but was still statistically different from the vehicle control. By day 84, there was no statistical difference between the sham mice with full dose CGRP with PZN and any of the other groups (Figure 5B).

As with concomitant administration of PZN, post-mTBI administration of PZN also resulted in faster recovery in males compared to females (Figure 5E-J). Males showed full recovery at day 42, while females fully recovered only at day 56. This sex difference was also seen with the PZN rescue of full dose CGRP in sham mice. There was a complete rescue in male, but not female mice at day 56 (Figure 5G, J).

Finally, identical to the concomitant PZN treatment, after PZN withdrawal, both groups experienced a significant increase in sensitivity compared to baseline, but these were restored within 2 weeks, by day 84, upon reintroducing PZN, with no sex differences (Figure 5B, E, H). Overall, these results show that continuous PZN treatment is effective when administered 2 weeks post-mTBI and confirm that PZN acts faster in male than female CD1 mice.

### CGRP disrupts glymphatic transport and chronic PZN restores glymphatic function

To understand why PZN was able to rescue CGRP-induced periorbital allodynia in mice that had not had mTBI, we examined the effect of CGRP on glymphatic function. Glymphatic transport was measured by injecting fluorescent Texas Red-conjugated dextran tracer into the cisterna magna of anaesthetized mice, as previously described^10^. Penetration of the tracer into the brain parenchyma was assessed by imaging brain slices along the rostro-caudal axis. CD1 mice were given drinking water with or without PZN for 6 weeks, then treated with full dose CGRP (0.1 mg kg^−1^) or vehicle ∼20 min before Texas Red-dextran infusion (Figure 6A). Three coronal brain slices were selected from the rostral portion of the brain and segmented into four regions of interest that correspond to sections 1, 3, and 5 as defined in a previous publication^40^ (Figure 6B). Three coronal brain slices were selected from the caudal portion of the brain and segmented into two regions of interest. Representative rostral and caudal brain sections are shown for each treatment (Figure 6C).

**Figure 6.**
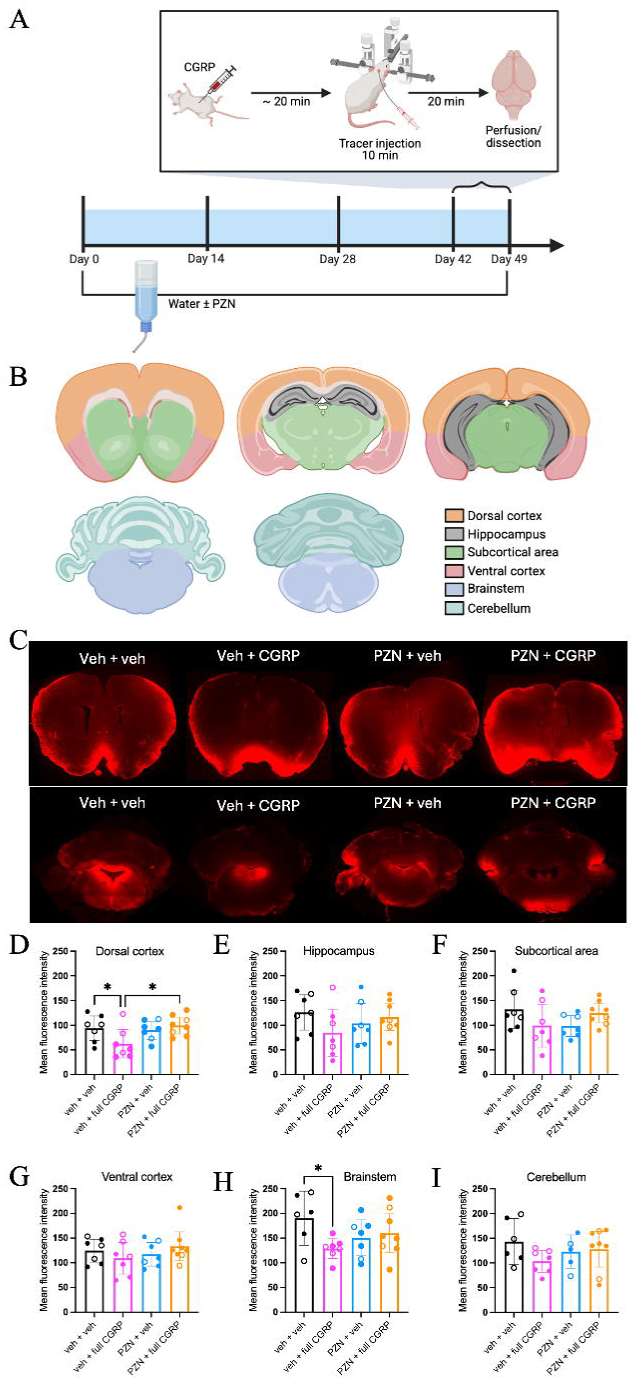
CGRP lowers and PZN restores glymphatic tracer influx in the dorsal cortex. **(A)** CD1 mice received drinking water with or without PZN until surgeries performed between days 42–49. Mice were anesthetized with ketamine/xylazine and administered CGRP (0.1 mg/kg) or vehicle, then prepared for ∼20 min prior to tracer injection. Tracer was infused for 10 min, followed by 20 min circulation period. Mice were then perfused and dissected. **(B)** Schematic illustrating the rostro–caudal brain section regions of interest. **(C)** Representative images of slices from a rostral section (top row) and a caudal section (bottom row) of the brain for the different treatments at 6 weeks: vehicle treated mice drinking water, CGRP treated mice drinking water, vehicle treated mice drinking PZN, and CGRP treated mice drinking PZN. **(D)-(I)** Regional average fluorescence intensity of the regions of interest between the different treatment conditions. Experiments performed in a single cohort of mice. Statistics in Supplementary Table 1.

Mice injected with CGRP exhibited reduced glymphatic CSF tracer influx into the dorsal cortex compared to mice treated with vehicle (Figure 6D). Mice given PZN for 6 weeks prior to CGRP injection exhibited glymphatic CSF tracer influx comparable to control mice given vehicle (Figure 6D). PZN treatment alone did not alter glymphatic transport in the dorsal cortex compared with vehicle-treated mice. There were no significant differences detected in other rostral brain regions (hippocampus, subcortical area, ventral cortex) (Figure 6E-G). In the brainstem, CGRP reduced glymphatic CSF tracer influx (Figure 6H). However, it is not clear if PZN could rescue this reduction since there was no statistical difference between the PZN and CGRP+PZN groups from either the vehicle control or CGRP groups (Figure 6H). In the cerebellum, glymphatic CSF tracer influx was not significantly affected by any of the treatments (Figure 6I). Thus, these data show that there is a CGRP-induced glymphatic impairment in the dorsal cortex and brainstem and that PZN can fully rescue this reduction in the dorsal cortex.

### *Aqp4* gene deletion sensitizes mice to CGRP-induced periorbital allodynia in the absence of mTBI

To test whether the PZN rescue of allodynia was indeed mediated by changes in glymphatic transport, we used AQP4 KO mice, in which glymphatic CSF influx and interstitial solute efflux are impaired^10,22,42^. Wildtype (WT) controls were established from nontransgenic littermates and maintained by breeding with C57BL/6J mice in parallel cages with AQP4 KO mice.

We assessed the periorbital sensitivity of AQP4 KO mice to CGRP. Mice were injected 30 min prior to testing with progressively lower doses of CGRP (Figure 7A). Among all the tested doses, as expected, WT mice only respond to CGRP 0.01 mg kg^−1^ and 0.5 mg kg^−1^. Instead, AQP4 KO mice showed significantly increased periorbital sensitivity at 0.01 mg kg^−1^ and 0.005 mg kg^−1^ compared to vehicle and to WT mice, but no longer displayed increased sensitivity at 0.001 mg kg^−1^ CGRP (Figure 7A). Therefore, the dose of 0.001 mg kg^−1^ of CGRP was used as the subthreshold dose and 0.01 mg kg^−1^ as the full dose for the AQP4 mice. To avoid terminology confusion when referring to the updated doses, all figures containing AQP4 KO mice will display the CGRP dose.

**Figure 7.**
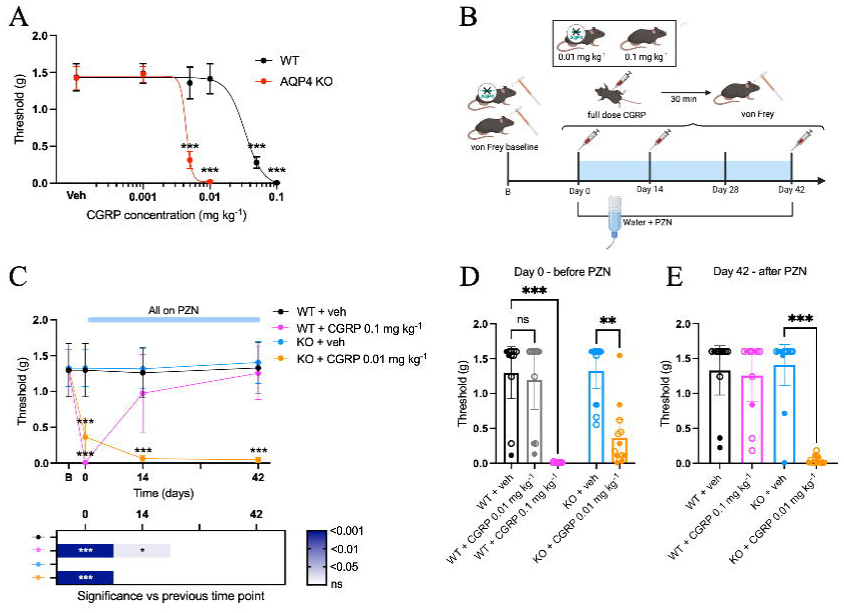
Lack of AQP4 channels sensitizes mice to CGRP-induced periorbital allodynia in the absence of mTBI. **(A)** Dose–response curve for CGRP induced periorbital sensitivity in AQP4 KO (red) and WT mice (black). All doses were tested in two independent cohorts, except for CGRP at 0.05 mg kg ¹, which was tested in only one cohort, and CGRP at 0.1 mg kg ¹, which was tested in a separate cohort. **(B)** Protocol schematic. Periorbital allodynia was assessed at baseline (B) and at the indicated time points 30 min after i.p. administration of vehicle, or CGRP (0.1 mg kg^−1^ for WT and 0.01 mg kg^−1^ for AQP4 KO mice). PZN treatment was started after the test indicated at day 0 in the drinking water of all the mice and continued for 6 weeks up to day 42. **(C)** Longitudinal data for all mice with significance between treatments: WT + CGRP 0.1 mg kg^−1^ vs WT + veh on one side, and KO + veh vs KO + CGRP 0.01 mg kg^−1^. The significance within each group for a time point compared to the previous one is reported in the heatmap. **(D)** Scatter plot of individual mice at day 0, before PZN administration, from the longitudinal plot. Open symbols for males and closed symbols for females. **(E)** Scatter plot of individual mice at day 42 from the longitudinal plot. Open symbols for males and closed symbols for females. Experiments from panels 7C-E performed in a single cohort of mice. Statistics in Supplementary Table 1.

We then tested whether PZN rescue of CGRP-induced allodynia in the absence of mTBI was sensitive to *Aqp4* gene deletion (Figure 7B). In contrast to WT mice, PZN was not able to rescue allodynia in the AQP4 KO mice (Figure 7C, E). AQP4 KO mice were sensitive to 0.01 mg kg^−1^ CGRP before PZN (day 0) and after 6 weeks PZN (day 42) (Figure D, E). Thus, the lack of AQP4 channels sensitized mice to CGRP and established that intact glymphatic machinery is required for PZN to be effective.

### PZN rescue of CGRP- and SNP-induced periorbital allodynia after mTBI requires AQP4 channels

To evaluate whether AQP4 channels are required for PZN rescue of allodynia following mTBI, we tested WT and AQP4 KO mice with PZN administered either after or concurrently with mTBI. For the post-mTBI PZN protocol, we used the newly identified subthreshold CGRP dose for AQP4 KO mice (0.001 mg kg ¹) (Figure 8A). In the absence of PZN, in the acute phase at day 1 after mTBI, both WT and KO mice had a significant increase in periorbital sensitivity compared to baseline (Figure 8B). At day 14, still in the absence of PZN, AQP4 KO, but not WT mice were sensitive to 0.001 mg kg ¹ (Figure 8B, C). At day 56, after 6 weeks of PZN treatment, CGRP-induced allodynia was not rescued by PZN treatment of AQP4 KO mice (Figure 8B, C).

**Figure 8.**
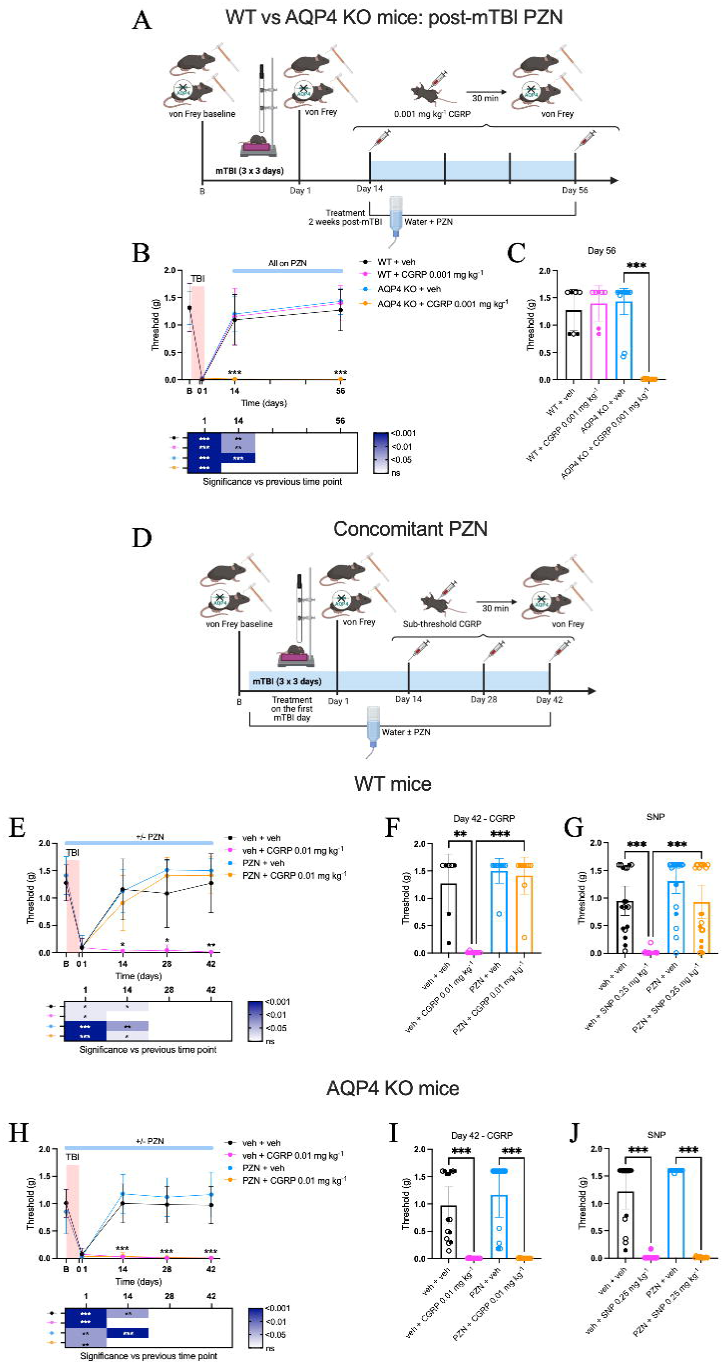
Rescue of CGRP and SNP sensitivity in mTBI WT but not in AQP4 KO mice with PZN treatment. **(A)** Protocol schematic. Periorbital allodynia was assessed at baseline (B) and at the indicated time points after mTBI without any treatment (day 1), then 30 min after i.p. administration of vehicle, or CGRP 0.001 mg kg^−1^ for AQP4 KO mice at day 14 and day 56. PZN treatment was started on day 14 post-mTBI in the drinking water of all the mice and continued for 6 weeks. **(B)** Longitudinal data for all WT vs AQP4 KO mTBI mice with significance between each treatment compared to veh + veh of their group reported on the graph. The significance within each group for a time point compared to the previous one is reported in the heatmap. **(C)** Scatter plot of individual mice at day 56 from the longitudinal plot. Open symbols for males and closed symbols for females. **(D)** Protocol schematic. Periorbital allodynia was assessed at baseline (B) and at the indicated time points after mTBI without any treatment (day1), and then 30 min after i.p. administration of vehicle, or CGRP 0.01 mg kg^−1^. PZN treatment was started on the first day of mTBI and continued for 6 weeks. **(E)** Longitudinal data for WT mice with significance between each treatment compared to veh + veh are reported on the graph. The significance within each group for a time point compared to the previous one is reported in the heatmap. **(F)** Scatter plot of individual WT mice at day 42 post-mTBI, showing single mice from the longitudinal data. Open symbols for males and closed symbols for females. **(G)** Scatter plot of WT mice at days 35 and 56 post mTBI, showing individual animals treated with SNP (0.25 mg/kg). Open symbols for males and closed symbols for females. This experiment was performed at later time points from the same mice of the longitudinal datasets shown in Figure 8E and at an intermediate time-point for Figure 2B. **(H)** Longitudinal data for AQP4 KO mice with significance between each treatment compared to veh + veh are reported on the graph. The significance within each group for a time point compared to the previous one is reported in the heatmap. **(I)** Scatter plot of individual AQP4 KO mice at day 42 post-mTBI from the longitudinal plot. Open symbols for males and closed symbols for females. **(J)** Scatter plot of AQP4 KO mice at days 35 and 56 post mTBI, showing individual animals treated with SNP (0.25 mg/kg). Open symbols for males and closed symbols for females. This experiment was performed at later time point from the same mice of longitudinal datasets shown in 8H. Experiments from panels 8B, C, E, F, H-J performed in a single cohort of WT and KO mice. Statistics in Supplementary Table 1.

For the concurrent mTBI PZN protocol, we used the subthreshold CGRP dose to match the dosage used in earlier WT experiments (0.01 mg kg ¹) (Figure 8D). Note that this dose is subthreshold for WT, but not for AQP4 KO mice. In the acute phase at day 1 after mTBI, both WT and KO mice had increased periorbital sensitivity compared to baseline (Figure 8E). Mice were then treated with CGRP every 2 weeks up to 42 days. As expected, CGRP-induced allodynia in WT mice was rescued by PZN (Figure 8E, F). In contrast, CGRP-induced allodynia was not rescued by PZN treatment of AQP4 KO mice (Figure 8H, I).

Finally, we tested a second migraine trigger, the nitric oxide donor sodium nitroprusside (SNP) at a subthreshold dosage for WT mice (0.25 mg kg^−1^)^39^. The same mice used for the CGRP trigger in Figure 8E, H and in Figure 2B were tested at days 56 and 35, respectively post-mTBI (Figure 8G). Similarly to CGRP, PZN fully rescued sensitivity to SNP induced by mTBI in WT mice (Figure 8G). Likewise, PZN did not rescue SNP-induced allodynia in AQP4 KO mice (Figure 8J).

## Discussion

This is the first study demonstrating that improved glymphatic function can alleviate tactile allodynia in migraine and PTH mouse models. We show that the migraine trigger CGRP reduces glymphatic transport and that directly contributes to allodynia. This mechanism likely dovetails with recent findings that CSF transport links cortical spreading depolarization to trigeminal nerve activation^27^, in addition to meningeal activation of the nerve^43,44^. The ability of the α1-noradrenergic antagonist PZN to restore glymphatic transport and alleviate periorbital allodynia provides a new mechanistic understanding of the pathogenesis of allodynia associated with migraine and PTH. Rescue by PZN was seen with both a full-dose CGRP-based migraine model and with two PTH models that used subthreshold CGRP and SNP doses. In addition, the phenotype was observed in two strains, outbred CD1 and inbred C57BL/6J mice. Evidence that PZN acts by enhancing glymphatic transport was shown by two approaches. First, fluorescence imaging of CSF tracer influx showed that PZN restored glymphatic transport that had been reduced by CGRP. Second, PZN was unable to rescue allodynia in mice lacking the AQP4 water channel required for glymphatic function. Furthermore, compared to WT, the AQP4 KO mice displayed an intrinsically heightened tactile sensitivity to a CGRP trigger. Together, these findings document that the mechanism of action of PZN rescue of migraine and PTH allodynia relies on glymphatic function.

The glymphatic system allows influx of cerebrospinal fluid into brain tissue and clearance of interstitial solutes out of brain tissue by perivascular pathways^10^. As such, glymphatic dysfunction has been proposed to contribute to an increasing number of neurological diseases, including migraine and TBI symptoms (for review^45^). Indeed, glymphatic function is profoundly impaired in mice following TBI^14,46^ (for reviews^11,17,47,48^). This glymphatic disruption has been associated with an abnormally high release of noradrenaline^46^. In a mouse model of moderate/severe TBI, perivascular CSF influx and interstitial solute efflux was impaired for at least 28 days following injury, in part due to the loss of perivascular AQP4 localization following TBI^14^. Several studies have reported disrupted glymphatic flow in migraine and PTH models^15,16,49,50^. Similar findings have been published for human MRI findings of enlargement of the perivascular spaces, which is commonly interpreted as an indirect marker of impaired perivascular function. An association between increased perivascular space volume and prior TBI has also been shown^18^. Overall, it remains an open question whether migraine pathology impairs the glymphatic system or altered glymphatic function contributes to migraine.

To address the cause and effect question, we built on a report showing that acute administration of a cocktail of noradrenergic receptor antagonists PZN, PPL, and atipamezole increased glymphatic transport^34^. Our results have narrowed that cocktail down to the α1-antagonist PZN and ruled out the β-antagonist PPL, although the role of the α2 antagonist atipamezole remains open. Importantly, we linked the effect of PZN on glymphatic function to tactile allodynia associated with migraine and PTH in mice by showing the dependence on AQP4 water channels. Thus, PZN can alleviate periorbital allodynia by rescuing glymphatic dysfunction.

A key finding of our study is that the migraine trigger CGRP reduces glymphatic transport in mice. This is consistent with a previous report that chronic treatment for 9 days with nitroglycerin, another known migraine trigger, reduced glymphatic function in the medullary dorsal horn in mice (only region reported)^51^. Following CGRP administration, we observed reduced glymphatic transport that was most pronounced in the dorsal cortex and brainstem. Notably, chronic PZN treatment could rescue the CGRP-reduced glymphatic CSF tracer influx in the dorsal cortex. Treatment with PZN alone did not apparently increase influx above vehicle levels. This latter point differs from the effect of a noradrenergic antagonist cocktail containing PZN in a prior study using awake, non-anesthetized mice^34^. This difference is likely due to awake mice having tonic noradrendergic output since anesthesia is known to suppress noradrenergic tone^22,34,52^. However, a limitation of this CSF tracer imaging method is that it only provides a static measurement. Therefore, we cannot exclude the possibility that more sensitive dynamic imaging approaches^40^ may reveal an effect of PZN alone and reveal changes in additional brain regions not seen in this study. Neverthless, region-dependent changes in glymphatic transport have been reported in another disorder, frontotemporal dementia^53^. Additionally, it has to be considered that the glymphatic transport is structurally dependent on the volume of the perivascular spaces and the polarization of AQP4 channels, features that may vary between brain regions (for review^54^). Coupled with the proposed roles of CGRP in migraine involving cerebrovasodilation^55^ and meningeal lymphatic vessel constriction^56^, our results squarely place CGRP at the intersection of cerebrovascular, lymphatic and glymphatic mechanisms contributing to headache.

What are the mechanisms by which CGRP reduces glymphatic function and PZN rescues this impairment? One likely mechanism may involve the vascular tone. CGRP gained early notoriety as the most potent vasodilatory peptide^57^, especially in the cerebrovasculature^58^. Human infusion studies confirmed that CGRP is a potent dilatator of cerebral arteries^59^. Importantly, cerebral vasodilation impairs glymphatic transport by closing off perivascular pathways^60^. Hence, CGRP reduction of glymphatic influx is consistent with its vasodilatory actions. However, a complexity is that the site of CGRP action on ex vivo perfused middle cerebral arteries is abluminal, not luminal^61^. Consequently, peripherally delivered CGRP in animal and human studies is likely to dilate cerebral arteries primarily near the surface of the brain where they enter the pia. Interestingly, CGRP only had a significant effect on tracer influx in the dorsal cortical and brainstem regions and not in deeper regions. However, deeper vasodilatory effects of CGRP could still occur indirectly by an endothelial nitric oxide mechanism (for review^62^). This indirect mechanism is apparently less robust than abluminal dilation since luminal CGRP-mediated dilation was statistically insignificant in the perfusion study^61^. PZN would thus restore the impaired influx primarily by increasing the interstitial volume fraction, in a manner similar to that reported for the cocktail of noradrenergic receptor antagonists^34^.

A second mechanism could involve CGRP activation of central noradrenergic signaling. One of the earliest CGRP studies reported that intracerebroventricular injection of CGRP caused central norepinephrine release^63^. Subsequent studies reported that peripheral CGRP, while vasodilatory, also caused a marked sympathetic outflow^64^. Furthermore, CGRP is associated with pain responses^65^ and peripheral injection of CGRP causes spontaneous grimace^66^. Indeed, activation of the pain matrix leads to both central and peripheral sympathetic activity. Noradrenergic signaling suppresses glymphatic function^67^, and this disruption may be reversed by PZN antagonism.

Finally, it is conceptually possible that CGRP could affect AQP4 localization, as suggested after mTBI^31^, which might then be modulated by PZN. However, given the rapid effect of CGRP to reduce glymphatic function, downregulation of AQP4 expression does not seem likely, as suggested following chronic nitroglycerin treatment^51^. Future studies are required to identify the mechanisms involved in interstitial space modulation during noradrenergic receptor blockade. Notably, this mechanism should be selective for α1 receptors, as β-noradrenergic receptor antagonism was ineffective in our mTBI mouse model.

In our mouse models, the modality and concentration of PZN administration was a critical parameter. High acute levels of PZN induced allodynia in mice, possibly due to the transient vasodilatory and hypotensive effects of PZN^68^. While lower acute concentrations did not produce detectable allodynia or non-evoked pain, they also lacked therapeutic efficacy. These observations underline the need for chronic PZN treatment.

Unexpectedly, the β-noradrenergic antagonist PPL was not effective in rescuing CGRP-induced allodynia in our PTH mouse model. This was surprising since PPL is used for migraine prevention and the European Headache Federation has recommended PPL as a first line treatment^38^. However, despite this designation, PPL has shown only limited efficacy in some clinical trials^69,70^. Overall, these results indicate that, while both are noradrenergic antagonists, PPL does not work by the same mechanism as PZN.

The possibility that α1-noradrenergic antagonism might be clinically beneficial for migraine was actually suggested over 30 years ago. A small (n=20) single blind, placebo-controlled study reported that thymoxamine, an α1-noradrenergic antagonist, was able to abort nitroglycerin-induced headache^71^. The same group also suggested that the antimigraine effect of ergotamine might be due to α1-noradrenergic antagonism activity^71^, although the complexity of ergotamine actions at multiple receptors precluded any firm conclusion. Interestingly, in a small nonblinded and uncontrolled series of observations, prophylactic use of an α1-noradrenergic antagonist, either terazosin or doxazosin (quinazoline derivatives similar to PZN), reduced the duration and frequency of headache in 10 patients^72^. Interestingly, more recently terazosin has also been shown to reduce Parkinson’s and Alzheimer’s progression^73–76^. However, the mechanism of action in these disorders was not α1-antagonism, but rather a metabolic action of terazosin that increased the activity of phosphoglycerate kinase 1, thereby increasing cellular ATP levels. To our knowledge, PZN has not been shown to have this metabolic activity. While speculative, it will be interesting to see if terazosin may be even more efficacious than PZN for treating migraine since mitochondrial dysfunction and metabolic alterations have been suggested to occur in migraine (for review^77^).

The best evidence for PZN as a PTH therapeutic is from a randomized, controlled trial of PZN administered for 12 weeks to Veterans and active-duty soldiers^36^. There was a significantly greater reduction of headache days for PZN- versus placebo-treated participants. This small clinical study suggests that PZN can be effective in the treatment of PTH, and along with our rodent data supports the hypothesis that glymphatic impairment following mTBI may contribute to the development of PTH-associated symptoms.

Finally, a significant advantage of PZN as a therapeutic is that it is relatively safe and has been on the market for about 50 years, and is still used by ∼500,000 patients yearly^78^. PZN was initially developed to treat high blood pressure but has been supplanted by newer drugs (for review^79^). More recently it has been used to treat benign enlargement of the prostate (for review^80^) and to reduce PTSD nightmares (for review^81^). While its efficacy for PTSD was shown by several trials^82–86^, this has been called into question by a large multisite clinical trial reporting that combat Veterans did no better with PZN than placebo^87^. Intriguingly, there were subgroups of Veterans who did benefit from PZN. These responders reported hypersensitivity described as an “adrenaline storm”^88^. This description is consistent with the pharmacology of PZN as a noradrenaline blocker. While the mechanism by which PZN improves nightmares is not known, it is tempting to speculate that reduced noradrenergic activity during REM sleep improves glymphatic removal of toxic and inflammatory agents from the brain.

Overall, our study proposes α1-noradrenergic receptor antagonism as a novel therapeutic strategy for allodynia in both migraine and PTH. We demonstrate that chronic PZN treatment alleviates periorbital tactile allodynia across migraine and PTH mouse models and strains and provides genetic evidence that its efficacy depends on intact glymphatic function. Our work links CGRP-induced headache pain to glymphatic impairment and shows that this dysfunction can be restored by PZN. These findings offer a mechanistic explanation for prior clinical observations and supports the rationale for optimizing α1-noradrenergic antagonists, particularly dosing and treatment paradigms, for future translational and clinical studies targeting both migraine and PTH patients.

## Data availability

The data that support the findings of this study are available from the corresponding author upon reasonable request.

## Acknowledgments

We thank Dr. Anne-Sophie Wattiez for establishing the mTBI protocol and initiating the prazosin and mTBI experiments. Protocol schematics were designed with Biorender.com.

## Supplementary material

Supplementary material is available at Brain online.

