## Supplementary Table 1 for "Glymphatic function restored by *α*1-noradrenergic antagonism alleviates headache allodynia in mice"

### Figure 1B

Mixed effect analysis

| | |
|---|---|
| Number of families | 11 |
| Number of comparisons per row family | 6 |
| Number of comparisons per column fami | 21 |
| Alpha | 0.05 |

| Fixed effects (type III) | P value | P value summary | Statistically significant (P < 0.05)? | F (DFn, DFd) | Geisser-Greenhouse's epsilon |
|---|---|---|---|---|---|
| Time | <.001 | *** | Yes | F (4.423, 459.9) = 37.72 | 0.7371 |
| Treatment Factor | <.001 | *** | Yes | F (3, 137) = 52.06 | |
| Time x Treatment Factor | <.001 | *** | Yes | F (13.27, 459.9) = 11.18 | 0.7371 |

| Tukey's multiple comparisons test | Mean diff. | 95.00% CI of diff. | Summary | Adjusted P Value | n1 | n2 |
|---|---|---|---|---|---|---|
| B | | | | | | |
| PZN + veh vs. veh + veh | -0.1043 | -0.6474 to 0.4387 | ns | 0.957 | 35 | 35 |
| PZN + subCGRP vs. veh + veh | -0.1227 | -0.6664 to 0.4210 | ns | 0.934 | 36 | 35 |
| veh + veh vs. veh + subCGRP | 0.08617 | -0.4376 to 0.6100 | ns | 0.973 | 35 | 35 |
| Day 1 | | | | | | |
| PZN + veh vs. veh + veh | -0.0359 | -0.3135 to 0.2417 | ns | 0.986 | 35 | 35 |
| PZN + subCGRP vs. veh + veh | 0.008867 | -0.3110 to 0.3287 | ns | >.999 | 36 | 35 |
| veh + veh vs. veh + subCGRP | -0.01119 | -0.3320 to 0.3096 | ns | >.999 | 35 | 35 |
| Day 14 | | | | | | |
| PZN + veh vs. veh + veh | -0.01577 | -0.4929 to 0.4614 | ns | >.999 | 22 | 23 |
| PZN + subCGRP vs. veh + veh | -0.7104 | -1.219 to -0.2020 | ** | 0.003 | 24 | 23 |
| veh + veh vs. veh + subCGRP | 1.118 | 0.6615 to 1.575 | *** | <.001 | 23 | 23 |
| Day 28 | | | | | | |
| PZN + veh vs. veh + veh | 0.005776 | -0.5770 to 0.5886 | ns | >.999 | 23 | 23 |
| PZN + subCGRP vs. veh + veh | -0.4672 | -1.044 to 0.1095 | ns | 0.15 | 24 | 23 |
| veh + veh vs. veh + subCGRP | 1.202 | 0.7646 to 1.640 | *** | <.001 | 23 | 23 |
| Day 42 | | | | | | |
| PZN + veh vs. veh + veh | -0.02962 | -0.4090 to 0.3498 | ns | 0.997 | 35 | 35 |
| PZN + subCGRP vs. veh + veh | -0.2357 | -0.6284 to 0.1569 | ns | 0.396 | 36 | 35 |
| veh + veh vs. veh + subCGRP | 1.381 | 1.082 to 1.680 | *** | <.001 | 35 | 35 |
| Day 56 | | | | | | |
| PZN + veh vs. veh + veh | 0.1936 | -0.1273 to 0.5144 | ns | 0.381 | 22 | 23 |
| PZN + subCGRP vs. veh + veh | -1.201 | -1.472 to -0.9295 | *** | <.001 | 24 | 23 |
| veh + veh vs. veh + subCGRP | 1.195 | 0.9202 to 1.469 | *** | <.001 | 23 | 22 |
| Day 84 | | | | | | |
| PZN + veh vs. veh + veh | 0.09744 | -0.2959 to 0.4908 | ns | 0.91 | 23 | 22 |
| PZN + subCGRP vs. veh + veh | 0.03471 | -0.3995 to 0.4689 | ns | 0.996 | 22 | 22 |
| veh + veh vs. veh + subCGRP | 1.039 | 0.6383 to 1.439 | *** | <.001 | 22 | 23 |
| veh + veh | | | | | | |
| B vs. day 1 | 0.9039 | 0.4460 to 1.362 | *** | <.001 | 35 | 35 |
| Day 1 vs. day 14 | -1.084 | -1.587 to -0.5810 | *** | <.001 | 23 | 23 |
| Day 14 vs. day 28 | -0.07678 | -0.6239 to 0.4703 | ns | >.999 | 23 | 23 |
| Day 28 vs. day 42 | -0.2187 | -0.8473 to 0.4098 | ns | 0.913 | 23 | 23 |
| Day 42 vs. day 56 | 0.01773 | -0.3196 to 0.3551 | ns | >.999 | 23 | 23 |
| Day 56 vs. day 84 | 0.05758 | -0.4178 to 0.5329 | ns | >.999 | 22 | 22 |
| veh + subCGRP | | | | | | |
| B vs. day 1 | 0.8065 | 0.3626 to 1.250 | *** | <.001 | 35 | 35 |
| Day 1 vs. day 14 | -0.01269 | -0.5960 to 0.5707 | ns | >.999 | 23 | 23 |
| Day 14 vs. day 28 | 0.007091 | -0.4742 to 0.4884 | ns | >.999 | 23 | 23 |
| Day 28 vs. day 42 | 0.1702 | -0.1287 to 0.4690 | ns | 0.535 | 23 | 23 |
| Day 42 vs. day 56 | 0.02936 | -0.2249 to 0.2836 | ns | >.999 | 22 | 22 |
| Day 56 vs. day 84 | -0.13 | -0.4175 to 0.1574 | ns | 0.758 | 22 | 22 |
| PZN + veh | | | | | | |
| B vs. day 1 | 0.8354 | 0.3865 to 1.284 | *** | <.001 | 35 | 35 |
| Day 1 vs. day 14 | -1.12 | -1.511 to -0.7286 | *** | <.001 | 22 | 22 |
| Day 14 vs. day 28 | -0.09408 | -0.6083 to 0.4201 | ns | 0.996 | 22 | 22 |
| Day 28 vs. day 42 | -0.08399 | -0.6005 to 0.4325 | ns | 0.998 | 23 | 23 |
| Day 42 vs. day 56 | -0.09641 | -0.5083 to 0.3154 | ns | 0.986 | 22 | 22 |
| Day 56 vs. day 84 | 0.1208 | -0.2920 to 0.5337 | ns | 0.959 | 22 | 22 |
| PZN + subCGRP | | | | | | |
| B vs. day 1 | 0.7724 | 0.3252 to 1.220 | *** | <.001 | 36 | 36 |
| Day 1 vs. day 14 | -0.4045 | -0.7777 to -0.03119 | * | 0.028 | 24 | 24 |
| Day 14 vs. day 28 | -0.3199 | -0.8207 to 0.1809 | ns | 0.408 | 24 | 24 |
| Day 28 vs. day 42 | -0.2248 | -0.8881 to 0.4385 | ns | 0.924 | 24 | 24 |
| Day 42 vs. day 56 | 1.256 | 0.9483 to 1.564 | *** | <.001 | 24 | 24 |
| Day 56 vs. day 84 | -1.202 | -1.617 to -0.7864 | *** | <.001 | 22 | 22 |

### Figure 1C

Kruskal-Wallis test

| | |
|---|---|
| Number of families | 1 |
| Number of comparisons per family | 3 |
| Alpha | 0.05 |

| | |
|---|---|
| P value | <.001 |
| Exact or approximate P value? | Approximate |
| P value summary | *** |
| Do the medians vary signif. (P < 0.05)? | Yes |
| Number of groups | 4 |
| Kruskal-Wallis statistic | 71.52 |

| Dunn's multiple comparisons test | Mean rank diff. | Significant? | Summary | Adjusted P Value | n1 | n2 |
|---|---|---|---|---|---|---|
| veh + veh vs. veh + subCGRP | 69.87 | Yes | *** | <.001 | 35 | 35 |
| veh + subCGRP vs. PZN + subCGRP | -57.3 | Yes | *** | <.001 | 35 | 36 |
| PZN + veh vs. PZN + subCGRP | 10.09 | No | ns | 0.859 | 35 | 36 |

### Figure 1D

Mixed effect analysis

| | |
|---|---|
| Number of families | 11 |
| Number of comparisons per row family | 6 |
| Number of comparisons per column fami | 21 |
| Alpha | 0.05 |

| Fixed effects (type III) | P value | P value summary | Statistically significant (P < 0.05)? | F (DFn, DFd) | Geisser-Greenhouse's epsilon |
|---|---|---|---|---|---|
| Time | <.001 | *** | Yes | F (3.751, 185.0) = 21.36 | 0.6251 |
| Treatment Factor | <.001 | *** | Yes | F (3, 66) = 21.92 | |
| Time x Treatment Factor | <.001 | *** | Yes | F (11.25, 185.0) = 5.940 | 0.6251 |

| Tukey's multiple comparisons test | Mean diff. | 95.00% CI of diff. | Summary | Adjusted P Value | n1 | n2 |
|---|---|---|---|---|---|---|
| B | | | | | | |
| PZN + veh vs. veh + veh | -0.04969 | -0.9435 to 0.8441 | ns | 0.999 | 18 | 17 |
| PZN + subCGRP vs. veh + veh | -0.01799 | -0.8884 to 0.8525 | ns | >.999 | 18 | 17 |
| veh + veh vs. veh + subCGRP | 0.0353 | -0.7871 to 0.8577 | ns | >.999 | 17 | 17 |
| Day 1 | | | | | | |
| PZN + veh vs. veh + veh | -0.02356 | -0.4170 to 0.3699 | ns | 0.998 | 18 | 17 |
| PZN + subCGRP vs. veh + veh | 0.1083 | -0.4161 to 0.6327 | ns | 0.942 | 18 | 17 |
| veh + veh vs. veh + subCGRP | -0.108 | -0.6492 to 0.4331 | ns | 0.947 | 17 | 17 |
| Day 14 | | | | | | |
| PZN + veh vs. veh + veh | -0.1684 | -0.7561 to 0.4194 | ns | 0.852 | 11 | 11 |
| PZN + subCGRP vs. veh + veh | -0.5478 | -1.244 to 0.1482 | ns | 0.155 | 12 | 11 |
| veh + veh vs. veh + subCGRP | 1.431 | 0.9815 to 1.880 | *** | <.001 | 11 | 11 |
| Day 28 | | | | | | |
| PZN + veh vs. veh + veh | 0.08287 | -0.7581 to 0.9239 | ns | 0.992 | 12 | 11 |
| PZN + subCGRP vs. veh + veh | -0.09463 | -0.8244 to 0.6352 | ns | 0.983 | 12 | 11 |
| veh + veh vs. veh + subCGRP | 1.122 | 0.5241 to 1.719 | *** | <.001 | 11 | 11 |
| Day 42 | | | | | | |
| PZN + veh vs. veh + veh | -0.04773 | -0.6955 to 0.6001 | ns | 0.997 | 18 | 17 |
| PZN + subCGRP vs. veh + veh | -0.4616 | -1.115 to 0.1917 | ns | 0.243 | 18 | 17 |
| veh + veh vs. veh + subCGRP | 1.575 | 1.085 to 2.065 | *** | <.001 | 17 | 17 |
| Day 56 | | | | | | |
| PZN + veh vs. veh + veh | 0.2267 | -0.3483 to 0.8016 | ns | 0.691 | 12 | 11 |
| PZN + subCGRP vs. veh + veh | -1.083 | -1.570 to -0.5951 | *** | <.001 | 12 | 11 |
| veh + veh vs. veh + subCGRP | 1.057 | 0.5616 to 1.553 | *** | <.001 | 11 | 10 |
| Day 84 | | | | | | |
| PZN + veh vs. veh + veh | 0.1052 | -0.5074 to 0.7177 | ns | 0.962 | 12 | 10 |

| | | | | | | |
|---|---|---|---|---|---|---|
| PZN + subCGRP vs. veh + veh | 0.08813 | -0.5690 to 0.7453 | ns | 0.981 | 11 | 10 |
| veh + veh vs. veh + subCGRP | 1.02 | 0.3857 to 1.655 | ** | 0.001 | 10 | 11 |
| veh + veh | | | | | | |
| B vs. day 1 | 1.022 | 0.2599 to 1.784 | ** | 0.005 | 17 | 17 |
| Day 1 vs. day 14 | -1.213 | -2.008 to -0.4190 | ** | 0.003 | 11 | 11 |
| Day 14 vs. day 28 | 0.1353 | -0.7312 to 1.002 | ns | 0.997 | 11 | 11 |
| Day 28 vs. day 42 | -0.5928 | -1.466 to 0.2805 | ns | 0.268 | 11 | 11 |
| Day 42 vs. day 56 | 0.147 | -0.1131 to 0.4070 | ns | 0.444 | 11 | 11 |
| Day 56 vs. day 84 | -0.02415 | -0.7318 to 0.6835 | ns | >.999 | 10 | 10 |
| veh + subCGRP | | | | | | |
| B vs. day 1 | 0.8785 | 0.1108 to 1.646 | * | 0.02 | 17 | 17 |
| Day 1 vs. day 14 | 0.3428 | -0.6006 to 1.286 | ns | 0.83 | 11 | 11 |
| Day 14 vs. day 28 | -0.1739 | -0.4735 to 0.1256 | ns | 0.416 | 11 | 11 |
| Day 28 vs. day 42 | 0.1352 | -0.3516 to 0.6221 | ns | 0.941 | 11 | 11 |
| Day 42 vs. day 56 | -0.05905 | -0.2280 to 0.1099 | ns | 0.838 | 10 | 10 |
| Day 56 vs. day 84 | -0.1206 | -0.6131 to 0.3720 | ns | 0.962 | 10 | 10 |
| PZN + veh | | | | | | |
| B vs. day 1 | 0.9956 | 0.2318 to 1.759 | ** | 0.007 | 18 | 18 |
| Day 1 vs. day 14 | -1.189 | -1.619 to -0.7577 | *** | <.001 | 11 | 11 |
| Day 14 vs. day 28 | -0.1102 | -1.209 to 0.9886 | ns | >.999 | 11 | 11 |
| Day 28 vs. day 42 | -0.3453 | -0.9994 to 0.3087 | ns | 0.53 | 12 | 12 |
| Day 42 vs. day 56 | -0.08803 | -0.8632 to 0.6871 | ns | >.999 | 12 | 12 |
| Day 56 vs. day 84 | 0.04942 | -0.6865 to 0.7854 | ns | >.999 | 12 | 12 |
| PZN + subCGRP | | | | | | |
| B vs. day 1 | 0.8954 | 0.1898 to 1.601 | ** | 0.008 | 18 | 18 |
| Day 1 vs. day 14 | -0.6205 | -1.209 to -0.03204 | * | 0.037 | 12 | 12 |
| Day 14 vs. day 28 | -0.3179 | -0.9507 to 0.3149 | ns | 0.581 | 12 | 12 |
| Day 28 vs. day 42 | 0.04785 | -0.7872 to 0.8829 | ns | >.999 | 12 | 12 |
| Day 42 vs. day 56 | 1.08 | 0.4845 to 1.676 | *** | <.001 | 12 | 12 |
| Day 56 vs. day 84 | -1.233 | -1.867 to -0.5994 | *** | <.001 | 11 | 11 |

**Figure 1E**

Kruskal-Wallis test

| | |
|---|---|
| Number of families | 1 |
| Number of comparisons per family | 3 |
| Alpha | 0.05 |

| | |
|---|---|
| P value | <.001 |
| Exact or approximate P value? | Approximate |
| P value summary | *** |
| Do the medians vary signif. (P < 0.05)? | Yes |
| Number of groups | 4 |
| Kruskal-Wallis statistic | 36.99 |

| Dunn's multiple comparisons test | Mean rank diff. | Significant? | Summary | Adjusted P Value | n1 | n2 |
|---|---|---|---|---|---|---|
| veh + veh vs. veh + subCGRP | 36.44 | Yes | *** | <.001 | 17 | 17 |
| veh + subCGRP vs. PZN + subCGRP | -26.37 | Yes | *** | <.001 | 17 | 18 |
| PZN + veh vs. PZN + subCGRP | 8.694 | No | ns | 0.572 | 18 | 18 |

**Figure 1F**

Mixed effect analysis

| | |
|---|---|
| Number of families | 11 |
| Number of comparisons per row family | 6 |
| Number of comparisons per column fami | 21 |
| Alpha | 0.05 |

| Fixed effects (type III) | P value | P value summary | Statistically significant (P < 0.05)? | F (DFn, DFd) | Geisser-Greenhouse's epsilon |
|---|---|---|---|---|---|
| Time | <.001 | *** | Yes | F (4.222, 213.9) = 16.33 | 0.7036 |
| Treatment Factor | <.001 | *** | Yes | F (3, 67) = 36.58 | |
| Time x Treatment Factor | <.001 | *** | Yes | F (12.67, 213.9) = 6.236 | 0.7036 |

| Tukey's multiple comparisons test | Mean diff. | 95.00% CI of diff. | Summary | Adjusted P Value | n1 | n2 |
|---|---|---|---|---|---|---|
| B | | | | | | |
| PZN + veh vs. veh + veh | -0.171 | -0.8489 to 0.5069 | ns | 0.903 | 17 | 18 |
| PZN + subCGRP vs. veh + veh | -0.2316 | -0.9403 to 0.4770 | ns | 0.814 | 18 | 18 |
| veh + veh vs. veh + subCGRP | 0.1342 | -0.5794 to 0.8479 | ns | 0.957 | 18 | 18 |
| Day 1 | | | | | | |
| PZN + veh vs. veh + veh | -0.04432 | -0.4651 to 0.3765 | ns | 0.992 | 17 | 18 |
| PZN + subCGRP vs. veh + veh | -0.08834 | -0.4978 to 0.3211 | ns | 0.936 | 18 | 18 |
| veh + veh vs. veh + subCGRP | 0.08024 | -0.3243 to 0.4848 | ns | 0.949 | 18 | 18 |
| Day 14 | | | | | | |
| PZN + veh vs. veh + veh | 0.1231 | -0.6816 to 0.9278 | ns | 0.973 | 11 | 12 |
| PZN + subCGRP vs. veh + veh | -0.8867 | -1.594 to -0.1797 | * | 0.011 | 12 | 12 |
| veh + veh vs. veh + subCGRP | 0.8321 | 0.02628 to 1.638 | * | 0.041 | 12 | 12 |
| Day 28 | | | | | | |
| PZN + veh vs. veh + veh | -0.07008 | -0.9813 to 0.8411 | ns | 0.996 | 11 | 12 |
| PZN + subCGRP vs. veh + veh | -0.8358 | -1.737 to 0.06506 | ns | 0.075 | 12 | 12 |
| veh + veh vs. veh + subCGRP | 1.276 | 0.5620 to 1.991 | *** | <.001 | 12 | 12 |
| Day 42 | | | | | | |
| PZN + veh vs. veh + veh | -0.032 | -0.4158 to 0.3518 | ns | 0.996 | 17 | 18 |
| PZN + subCGRP vs. veh + veh | -0.02036 | -0.4901 to 0.4493 | ns | >.999 | 18 | 18 |
| veh + veh vs. veh + subCGRP | 1.198 | 0.8297 to 1.566 | *** | <.001 | 18 | 18 |
| Day 56 | | | | | | |
| PZN + veh vs. veh + veh | 0.1857 | -0.1430 to 0.5144 | ns | 0.369 | 10 | 12 |
| PZN + subCGRP vs. veh + veh | -1.309 | -1.646 to -0.9727 | *** | <.001 | 12 | 12 |
| veh + veh vs. veh + subCGRP | 1.319 | 0.9838 to 1.653 | *** | <.001 | 12 | 12 |
| Day 84 | | | | | | |
| PZN + veh vs. veh + veh | 0.09148 | -0.4872 to 0.6701 | ns | 0.97 | 11 | 12 |
| PZN + subCGRP vs. veh + veh | -0.01711 | -0.6741 to 0.6399 | ns | >.999 | 11 | 12 |
| veh + veh vs. veh + subCGRP | 1.055 | 0.4734 to 1.637 | *** | <.001 | 12 | 12 |
| veh + veh | | | | | | |
| B vs. day 1 | 0.7926 | 0.1638 to 1.421 | ** | 0.009 | 18 | 18 |
| Day 1 vs. day 14 | -0.9657 | -1.760 to -0.1711 | * | 0.015 | 12 | 12 |
| Day 14 vs. day 28 | -0.2712 | -1.103 to 0.5605 | ns | 0.896 | 12 | 12 |
| Day 28 vs. day 42 | 0.1242 | -0.8419 to 1.090 | ns | 0.999 | 12 | 12 |
| Day 42 vs. day 56 | -0.1007 | -0.7629 to 0.5614 | ns | 0.997 | 12 | 12 |
| Day 56 vs. day 84 | 0.1257 | -0.6620 to 0.9134 | ns | 0.997 | 12 | 12 |
| veh + subCGRP | | | | | | |
| B vs. day 1 | 0.7386 | 0.1531 to 1.324 | ** | 0.009 | 18 | 18 |
| Day 1 vs. day 14 | -0.3386 | -1.116 to 0.4393 | ns | 0.714 | 12 | 12 |
| Day 14 vs. day 28 | 0.173 | -0.7931 to 1.139 | ns | 0.994 | 12 | 12 |
| Day 28 vs. day 42 | 0.2022 | -0.2675 to 0.6719 | ns | 0.724 | 12 | 12 |
| Day 56 vs. day 84 | -0.138 | -0.5781 to 0.3022 | ns | 0.911 | 12 | 12 |
| PZN + veh | | | | | | |
| B vs. day 1 | 0.6658 | 0.1137 to 1.218 | * | 0.013 | 17 | 17 |
| Day 1 vs. day 14 | -1.052 | -1.827 to -0.2761 | ** | 0.008 | 11 | 11 |
| Day 14 vs. day 28 | -0.078 | -0.4923 to 0.3363 | ns | 0.991 | 11 | 11 |
| Day 28 vs. day 42 | 0.2011 | -0.6937 to 1.096 | ns | 0.978 | 11 | 11 |
| Day 42 vs. day 56 | -0.1065 | -0.5239 to 0.3109 | ns | 0.954 | 10 | 10 |
| Day 56 vs. day 84 | 0.2065 | -0.2972 to 0.7102 | ns | 0.729 | 10 | 10 |
| PZN + subCGRP | | | | | | |
| B vs. day 1 | 0.6493 | 0.005099 to 1.293 | * | 0.048 | 18 | 18 |
| Day 1 vs. day 14 | -0.1884 | -0.6936 to 0.3169 | ns | 0.827 | 12 | 12 |
| Day 14 vs. day 28 | -0.322 | -1.258 to 0.6138 | ns | 0.871 | 12 | 12 |
| Day 28 vs. day 42 | -0.4975 | -1.668 to 0.6733 | ns | 0.734 | 12 | 12 |
| Day 42 vs. day 56 | 1.432 | 1.195 to 1.670 | *** | <.001 | 12 | 12 |
| Day 56 vs. day 84 | -1.17 | -1.875 to -0.4661 | ** | 0.002 | 11 | 11 |

**Figure 1G**

Kruskal-Wallis test

| | |
|---|---|
| Number of families | 1 |
| Number of comparisons per family | 3 |
| Alpha | 0.05 |

| | |
|---|---|
| P value | <.001 |
| Exact or approximate P value? | Approximate |
| P value summary | *** |
| Do the medians vary signif. (P < 0.05)? | Yes |
| Number of groups | 4 |
| Kruskal-Wallis statistic | 34.61 |

| Dunn's multiple comparisons test | Mean rank diff. | Significant? | Summary | Adjusted P Value | n1 | n2 |
|---|---|---|---|---|---|---|
| veh + veh vs. veh + subCGRP | 33.25 | Yes | *** | <.001 | 18 | 18 |
| veh + subCGRP vs. PZN + subCGRP | -31.28 | Yes | *** | <.001 | 18 | 18 |
| PZN + veh vs. PZN + subCGRP | 0.2859 | No | ns | >.999 | 17 | 18 |

### Figure 2B

Two-way ANOVA

| | |
|---|---|
| Number of families | 11 |
| Number of comparisons per row family | 6 |
| Number of comparisons per column family | 21 |
| Alpha | 0.05 |

| Source of Variation | % of total variation | P value | P value summary | Significant? | Geisser-Greenhouse's epsilon |
|---|---|---|---|---|---|
| Time x Treatment Factor | 13.09 | <.001 | *** | Yes | 0.8907 |
| Time | 23.91 | <.001 | *** | Yes | 0.8907 |
| Treatment | 18.79 | <.001 | *** | Yes | |

| Tukey's multiple comparisons test | Mean diff. | 95.00% CI of diff. | Summary | Adjusted P Value | n1 | n2 |
|---|---|---|---|---|---|---|
| B | | | | | | |
| veh + veh vs. veh + subCGRP | 0.1361 | -0.3303 to 0.6026 | ns | 0.86 | 20 | 19 |
| veh + veh vs. PZN + veh | 0.2515 | -0.2220 to 0.7251 | ns | 0.49 | 20 | 20 |
| veh + veh vs. PZN + subCGRP | -0.0716 | -0.4994 to 0.3562 | ns | 0.969 | 20 | 20 |
| Day 1 | | | | | | |
| veh + veh vs. veh + subCGRP | -0.06507 | -0.2120 to 0.08189 | ns | 0.61 | 20 | 19 |
| veh + veh vs. PZN + veh | -0.01226 | -0.05768 to 0.03317 | ns | 0.887 | 20 | 20 |
| veh + veh vs. PZN + subCGRP | 0.0053 | -0.03318 to 0.04378 | ns | 0.982 | 20 | 20 |
| Day 14 | | | | | | |
| veh + veh vs. veh + subCGRP | 1.028 | 0.7394 to 1.316 | *** | <.001 | 20 | 19 |
| veh + veh vs. PZN + veh | 0.1233 | -0.2967 to 0.5434 | ns | 0.858 | 20 | 20 |
| veh + veh vs. PZN + subCGRP | 0.4069 | -0.05254 to 0.8664 | ns | 0.098 | 20 | 20 |
| Day 28 | | | | | | |
| veh + veh vs. veh + subCGRP | 1.111 | 0.7290 to 1.493 | *** | <.001 | 20 | 19 |
| veh + veh vs. PZN + veh | -0.1427 | -0.6553 to 0.3699 | ns | 0.877 | 20 | 20 |
| veh + veh vs. PZN + subCGRP | 0.07786 | -0.4390 to 0.5948 | ns | 0.977 | 20 | 20 |
| Day 42 | | | | | | |
| veh + veh vs. veh + subCGRP | 0.9884 | 0.6135 to 1.363 | *** | <.001 | 20 | 19 |
| veh + veh vs. PZN + veh | -0.1649 | -0.6499 to 0.3202 | ns | 0.798 | 20 | 20 |
| veh + veh vs. PZN + subCGRP | -0.006535 | -0.4677 to 0.4546 | ns | >.999 | 20 | 20 |
| Day 56 | | | | | | |
| veh + veh vs. veh + subCGRP | 1.047 | 0.6566 to 1.436 | *** | <.001 | 20 | 19 |
| veh + veh vs. PZN + veh | 0.1082 | -0.4118 to 0.6282 | ns | 0.943 | 20 | 20 |
| veh + veh vs. PZN + subCGRP | 0.9555 | 0.5510 to 1.360 | *** | <.001 | 20 | 20 |
| Day 70 | | | | | | |
| veh + veh vs. veh + subCGRP | 0.9497 | 0.5323 to 1.367 | *** | <.001 | 20 | 19 |
| veh + veh vs. PZN + veh | -0.01896 | -0.5273 to 0.4894 | ns | >.999 | 20 | 20 |
| veh + veh vs. PZN + subCGRP | -0.2861 | -0.7372 to 0.1649 | ns | 0.334 | 20 | 20 |
| veh + veh | | | | | | |
| B vs. day 1 | 1.251 | 0.8816 to 1.621 | *** | <.001 | 20 | 20 |
| Day 1 vs. day 14 | -1.084 | -1.393 to -0.7754 | *** | <.001 | 20 | 20 |
| Day 14 vs. day 28 | -0.02252 | -0.4552 to 0.4101 | ns | >.999 | 20 | 20 |
| Day 28 vs. day 42 | 0.07393 | -0.3585 to 0.5063 | ns | 0.997 | 20 | 20 |
| Day 42 vs. day 56 | -0.001445 | -0.4759 to 0.4730 | ns | >.999 | 20 | 20 |
| Day 56 vs. day 70 | 0.001705 | -0.5547 to 0.5581 | ns | >.999 | 20 | 20 |
| veh + subCGRP | | | | | | |
| B vs. day 1 | 1.05 | 0.6411 to 1.459 | *** | <.001 | 19 | 19 |
| Day 1 vs. day 14 | 0.0084 | -0.2273 to 0.2441 | ns | >.999 | 19 | 19 |
| Day 14 vs. day 28 | 0.06076 | -0.09782 to 0.2193 | ns | 0.858 | 19 | 19 |
| Day 28 vs. day 42 | -0.04851 | -0.2144 to 0.1174 | ns | 0.955 | 19 | 19 |
| Day 42 vs. day 56 | 0.05673 | -0.1073 to 0.2208 | ns | 0.906 | 19 | 19 |
| Day 56 vs. day 70 | -0.09517 | -0.3436 to 0.1533 | ns | 0.858 | 19 | 19 |
| PZN + veh | | | | | | |
| B vs. day 1 | 0.9877 | 0.5519 to 1.423 | *** | <.001 | 20 | 20 |
| Day 1 vs. day 14 | -0.9486 | -1.351 to -0.5467 | *** | <.001 | 20 | 20 |
| Day 14 vs. day 28 | -0.2886 | -0.8367 to 0.2596 | ns | 0.606 | 20 | 20 |
| Day 28 vs. day 42 | 0.05176 | -0.3430 to 0.4465 | ns | >.999 | 20 | 20 |
| Day 42 vs. day 56 | 0.2716 | -0.1843 to 0.7275 | ns | 0.471 | 20 | 20 |
| Day 56 vs. day 70 | -0.1255 | -0.5676 to 0.3167 | ns | 0.962 | 20 | 20 |
| PZN + subCGRP | | | | | | |
| B vs. day 1 | 1.328 | 0.9683 to 1.688 | *** | <.001 | 20 | 20 |
| Day 1 vs. day 14 | -0.6826 | -1.137 to -0.2282 | ** | 0.001 | 20 | 20 |
| Day 14 vs. day 28 | -0.3516 | -0.8532 to 0.1500 | ns | 0.293 | 20 | 20 |
| Day 28 vs. day 42 | -0.01047 | -0.5859 to 0.5650 | ns | >.999 | 20 | 20 |
| Day 42 vs. day 56 | 0.9606 | 0.5902 to 1.331 | *** | <.001 | 20 | 20 |
| Day 56 vs. day 70 | -1.24 | -1.576 to -0.9035 | *** | <.001 | 20 | 20 |

### Figure 2C

Kruskal-Wallis test

| | |
|---|---|
| Number of families | 1 |
| Number of comparisons per family | 3 |
| Alpha | 0.05 |

| | |
|---|---|
| P value | <.001 |
| Exact or approximate P value? | Approximate |
| P value summary | *** |
| Do the medians vary signif. (P < 0.05)? | Yes |
| Number of groups | 4 |
| Kruskal-Wallis statistic | 40.01 |

| Dunn's multiple comparisons test | Mean rank di | Significant? | Summary | Adjusted P Value | n1 | n2 |
|---|---|---|---|---|---|---|
| veh + veh vs. veh + subCGRP | 35.52 | Yes | *** | <.001 | 20 | 19 |
| veh + subCGRP vs. PZN + subCGRP | -36.29 | Yes | *** | <.001 | 19 | 20 |
| PZN + veh vs. PZN + subCGRP | 3.95 | No | ns | >.999 | 20 | 20 |

### Figure 2D

Two-way ANOVA

| | |
|---|---|
| Number of families | 11 |
| Number of comparisons per row family | 6 |
| Number of comparisons per column family | 21 |
| Alpha | 0.05 |

| Source of Variation | % of total variation | P value | P value summary | Significant? | Geisser-Greenhouse's epsilon |
|---|---|---|---|---|---|
| Time x Treatment Factor | 14.5 | <.001 | *** | Yes | 0.8781 |
| Time | 23.69 | <.001 | *** | Yes | 0.8781 |
| Treatment | 18.38 | <.001 | *** | Yes | |

| Tukey's multiple comparisons test | Mean diff. | 95.00% CI of diff. | Summary | Adjusted P Value | n1 | n2 |
|---|---|---|---|---|---|---|
| B | | | | | | |
| veh + veh vs. veh + subCGRP | 0.2815 | -0.3589 to 0.9218 | ns | 0.599 | 10 | 10 |
| veh + veh vs. PZN + veh | 0.5273 | -0.2010 to 1.256 | ns | 0.199 | 10 | 10 |
| veh + veh vs. PZN + subCGRP | 0.06113 | -0.4953 to 0.6176 | ns | 0.989 | 10 | 10 |
| Day 1 | | | | | | |
| veh + veh vs. veh + subCGRP | -0.09631 | -0.3962 to 0.2036 | ns | 0.754 | 10 | 10 |
| veh + veh vs. PZN + veh | -0.03012 | -0.1023 to 0.04206 | ns | 0.622 | 10 | 10 |
| veh + veh vs. PZN + subCGRP | -0.00827 | -0.06056 to 0.04402 | ns | 0.968 | 10 | 10 |
| Day 14 | | | | | | |
| veh + veh vs. veh + subCGRP | 1.031 | 0.5225 to 1.539 | *** | <.001 | 10 | 10 |
| veh + veh vs. PZN + veh | 0.02656 | -0.6094 to 0.6625 | ns | >.999 | 10 | 10 |
| veh + veh vs. PZN + subCGRP | 0.3523 | -0.3426 to 1.047 | ns | 0.493 | 10 | 10 |
| Day 28 | | | | | | |
| veh + veh vs. veh + subCGRP | 1.031 | 0.3519 to 1.711 | ** | 0.005 | 10 | 10 |
| veh + veh vs. PZN + veh | 0.06324 | -0.8584 to 0.9849 | ns | 0.997 | 10 | 10 |

| | | | | | | |
|---|---|---|---|---|---|---|
| veh + veh vs. PZN + subCGRP | -0.2315 | -0.9677 to 0.5046 | ns | 0.802 | 10 | 10 |
| Day 42 | | | | | | |
| veh + veh vs. veh + subCGRP | 1.034 | 0.4355 to 1.632 | ** | 0.001 | 10 | 10 |
| veh + veh vs. PZN + veh | 0.08473 | -0.7118 to 0.8813 | ns | 0.99 | 10 | 10 |
| veh + veh vs. PZN + subCGRP | 0.05978 | -0.6463 to 0.7658 | ns | 0.995 | 10 | 10 |
| Day 56 | | | | | | |
| veh + veh vs. veh + subCGRP | 1.074 | 0.4632 to 1.685 | ** | 0.002 | 10 | 10 |
| veh + veh vs. PZN + veh | 0.3494 | -0.4655 to 1.164 | ns | 0.627 | 10 | 10 |
| veh + veh vs. PZN + subCGRP | 1.003 | 0.3877 to 1.617 | ** | 0.003 | 10 | 10 |
| Day 70 | | | | | | |
| veh + veh vs. veh + subCGRP | 1.012 | 0.4103 to 1.614 | ** | 0.002 | 10 | 10 |
| veh + veh vs. PZN + veh | 0.1038 | -0.7263 to 0.9339 | ns | 0.984 | 10 | 10 |
| veh + veh vs. PZN + subCGRP | -0.4318 | -1.089 to 0.2257 | ns | 0.273 | 10 | 10 |
| veh + veh | | | | | | |
| B vs. day 1 | 1.405 | 0.9492 to 1.861 | *** | <.001 | 10 | 10 |
| Day 1 vs. day 14 | -1.112 | -1.670 to -0.5548 | *** | <.001 | 10 | 10 |
| Day 14 vs. day 28 | 0.0666 | -0.6006 to 0.7338 | ns | >.999 | 10 | 10 |
| Day 28 vs. day 42 | -0.08864 | -0.6762 to 0.4989 | ns | 0.997 | 10 | 10 |
| Day 42 vs. day 56 | 0.07296 | -0.4801 to 0.6260 | ns | 0.998 | 10 | 10 |
| Day 56 vs. day 70 | 0.04835 | -0.9687 to 1.065 | ns | >.999 | 10 | 10 |
| veh + subCGRP | | | | | | |
| B vs. day 1 | 1.027 | 0.3589 to 1.696 | ** | 0.004 | 10 | 10 |
| Day 1 vs. day 14 | 0.015 | -0.4967 to 0.5267 | ns | >.999 | 10 | 10 |
| Day 14 vs. day 28 | 0.06727 | -0.2716 to 0.4061 | ns | 0.986 | 10 | 10 |
| Day 28 vs. day 42 | -0.08647 | -0.4425 to 0.2696 | ns | 0.963 | 10 | 10 |
| Day 42 vs. day 56 | 0.1133 | -0.2280 to 0.4546 | ns | 0.866 | 10 | 10 |
| Day 56 vs. day 70 | -0.01349 | -0.06607 to 0.03909 | ns | 0.953 | 10 | 10 |
| PZN + veh | | | | | | |
| B vs. day 1 | 0.8476 | 0.06966 to 1.626 | * | 0.031 | 10 | 10 |
| Day 1 vs. day 14 | -1.055 | -1.652 to -0.4593 | ** | 0.001 | 10 | 10 |
| Day 14 vs. day 28 | 0.1033 | -0.7314 to 0.9379 | ns | 0.999 | 10 | 10 |
| Day 28 vs. day 42 | -0.06715 | -0.7643 to 0.6300 | ns | >.999 | 10 | 10 |
| Day 42 vs. day 56 | 0.3377 | -0.1989 to 0.8742 | ns | 0.323 | 10 | 10 |
| Day 56 vs. day 70 | -0.1973 | -0.8380 to 0.4435 | ns | 0.899 | 10 | 10 |
| PZN + subCGRP | | | | | | |
| B vs. day 1 | 1.336 | 0.7786 to 1.893 | *** | <.001 | 10 | 10 |
| Day 1 vs. day 14 | -0.7515 | -1.447 to -0.05567 | * | 0.033 | 10 | 10 |
| Day 14 vs. day 28 | -0.5173 | -1.271 to 0.2365 | ns | 0.247 | 10 | 10 |
| Day 28 vs. day 42 | 0.2027 | -0.6186 to 1.024 | ns | 0.961 | 10 | 10 |
| Day 42 vs. day 56 | 1.016 | 0.3306 to 1.701 | ** | 0.005 | 10 | 10 |
| Day 56 vs. day 70 | -1.386 | -1.847 to -0.9250 | *** | <.001 | 10 | 10 |

### Figure 2E

Kruskal-Wallis test

| | |
|---|---|
| Number of families | 1 |
| Number of comparisons per family | 3 |
| Alpha | 0.05 |

| | |
|---|---|
| P value | <.001 |
| Exact or approximate P value? | Approximate |
| P value summary | *** |
| Do the medians vary signif. (P < 0.05)? | Yes |
| Number of groups | 4 |
| Kruskal-Wallis statistic | 17.82 |

| Dunn's multiple comparisons test | Mean rank di | Significant? | Summary | Adjusted P Value | n1 | n2 |
|---|---|---|---|---|---|---|
| veh + veh vs. veh + subCGRP | 18.05 | Yes | ** | 0.001 | 10 | 10 |
| veh + subCGRP vs. PZN + subCGRP | -17.7 | Yes | ** | 0.002 | 10 | 10 |
| PZN + veh vs. PZN + subCGRP | -0.45 | No | ns | >.999 | 10 | 10 |

### Figure 2F

Two-way ANOVA

| | |
|---|---|
| Number of families | 11 |
| Number of comparisons per row family | 6 |
| Number of comparisons per column family | 21 |
| Alpha | 0.05 |

| Source of Variation | % of total variation | P value | P value summary | Significant? | Geisser-Greenhouse's epsilon |
|---|---|---|---|---|---|
| Time x Treatment Factor | 14.22 | <.001 | *** | Yes | 0.817 |
| Time | 24.71 | <.001 | *** | Yes | 0.817 |
| Treatment | 21.19 | <.001 | *** | Yes | |

| Tukey's multiple comparisons test | Mean diff. | 95.00% CI of diff. | Summary | Adjusted P Value | n1 | n2 |
|---|---|---|---|---|---|---|
| B | | | | | | |
| veh + veh vs. veh + subCGRP | -0.008559 | -0.7725 to 0.7554 | ns | >.999 | 10 | 9 |
| veh + veh vs. PZN + veh | -0.02422 | -0.7157 to 0.6672 | ns | >.999 | 10 | 10 |
| veh + veh vs. PZN + subCGRP | -0.2043 | -0.9175 to 0.5088 | ns | 0.849 | 10 | 10 |
| Day 1 | | | | | | |
| veh + veh vs. veh + subCGRP | -0.03064 | -0.1275 to 0.06621 | ns | 0.802 | 10 | 9 |
| veh + veh vs. PZN + veh | 0.00561 | -0.06376 to 0.07498 | ns | 0.995 | 10 | 10 |
| veh + veh vs. PZN + subCGRP | 0.01887 | -0.04881 to 0.08655 | ns | 0.831 | 10 | 10 |
| Day 14 | | | | | | |
| veh + veh vs. veh + subCGRP | 1.027 | 0.6474 to 1.406 | *** | <.001 | 10 | 9 |
| veh + veh vs. PZN + veh | 0.2201 | -0.4125 to 0.8527 | ns | 0.752 | 10 | 10 |
| veh + veh vs. PZN + subCGRP | 0.4615 | -0.2520 to 1.175 | ns | 0.281 | 10 | 10 |
| Day 28 | | | | | | |
| veh + veh vs. veh + subCGRP | 1.192 | 0.6574 to 1.726 | *** | <.001 | 10 | 9 |
| veh + veh vs. PZN + veh | -0.3486 | -0.8856 to 0.1884 | ns | 0.254 | 10 | 10 |
| veh + veh vs. PZN + subCGRP | 0.3872 | -0.4004 to 1.175 | ns | 0.518 | 10 | 10 |
| Day 42 | | | | | | |
| veh + veh vs. veh + subCGRP | 0.949 | 0.3855 to 1.512 | ** | 0.002 | 10 | 9 |
| veh + veh vs. PZN + veh | -0.4144 | -1.059 to 0.2305 | ns | 0.295 | 10 | 10 |
| veh + veh vs. PZN + subCGRP | -0.07285 | -0.7632 to 0.6175 | ns | 0.99 | 10 | 10 |
| Day 56 | | | | | | |
| veh + veh vs. veh + subCGRP | 1.019 | 0.3735 to 1.664 | ** | 0.004 | 10 | 9 |
| veh + veh vs. PZN + veh | -0.133 | -0.8585 to 0.5926 | ns | 0.952 | 10 | 10 |
| veh + veh vs. PZN + subCGRP | 0.9085 | 0.2454 to 1.572 | ** | 0.007 | 10 | 10 |
| Day 70 | | | | | | |
| veh + veh vs. veh + subCGRP | 0.8777 | 0.1600 to 1.595 | * | 0.014 | 10 | 9 |
| veh + veh vs. PZN + veh | -0.1417 | -0.8385 to 0.5551 | ns | 0.938 | 10 | 10 |
| veh + veh vs. PZN + subCGRP | -0.1404 | -0.8492 to 0.5684 | ns | 0.942 | 10 | 10 |
| veh + veh | | | | | | |
| B vs. day 1 | 1.098 | 0.4224 to 1.773 | ** | 0.002 | 10 | 10 |
| Day 1 vs. day 14 | -1.056 | -1.504 to -0.6088 | *** | <.001 | 10 | 10 |
| Day 14 vs. day 28 | -0.1116 | -0.8451 to 0.6219 | ns | 0.996 | 10 | 10 |
| Day 28 vs. day 42 | 0.2365 | -0.5253 to 0.9983 | ns | 0.896 | 10 | 10 |
| Day 42 vs. day 56 | -0.07585 | -1.019 to 0.8669 | ns | >.999 | 10 | 10 |
| Day 56 vs. day 70 | -0.04494 | -0.8355 to 0.7456 | ns | >.999 | 10 | 10 |
| veh + subCGRP | | | | | | |
| B vs. day 1 | 1.076 | 0.3923 to 1.759 | ** | 0.004 | 9 | 9 |
| Day 1 vs. day 14 | 0.001067 | -0.07125 to 0.07339 | ns | >.999 | 9 | 9 |
| Day 14 vs. day 28 | 0.05352 | -0.03093 to 0.1380 | ns | 0.3 | 9 | 9 |

| | | | | | | |
|---|---|---|---|---|---|---|
| Day 28 vs. day 42 | -0.006322 | -0.02827 to 0.01563 | ns | 0.912 | 9 | 9 |
| Day 42 vs. day 56 | -0.006133 | -0.05744 to 0.04518 | ns | 0.999 | 9 | 9 |
| Day 56 vs. day 70 | -0.1859 | -0.7853 to 0.4135 | ns | 0.882 | 9 | 9 |
| PZN + veh | | | | | | |
| B vs. day 1 | 1.128 | 0.5303 to 1.725 | *** | <.001 | 10 | 10 |
| Day 1 vs. day 14 | -0.8417 | -1.534 to -0.1495 | * | 0.017 | 10 | 10 |
| Day 14 vs. day 28 | -0.6804 | -1.352 to -0.008708 | * | 0.047 | 10 | 10 |
| Day 28 vs. day 42 | 0.1707 | -0.3856 to 0.7269 | ns | 0.901 | 10 | 10 |
| Day 42 vs. day 56 | 0.2056 | -0.6986 to 1.110 | ns | 0.973 | 10 | 10 |
| Day 56 vs. day 70 | -0.05365 | -0.8447 to 0.7374 | ns | >.999 | 10 | 10 |
| PZN + subCGRP | | | | | | |
| B vs. day 1 | 1.321 | 0.6987 to 1.944 | *** | <.001 | 10 | 10 |
| Day 1 vs. day 14 | -0.6136 | -1.396 to 0.1689 | ns | 0.153 | 10 | 10 |
| Day 14 vs. day 28 | -0.1859 | -1.024 to 0.6520 | ns | 0.976 | 10 | 10 |
| Day 28 vs. day 42 | -0.2236 | -1.208 to 0.7607 | ns | 0.973 | 10 | 10 |
| Day 42 vs. day 56 | 0.9055 | 0.3958 to 1.415 | ** | 0.001 | 10 | 10 |
| Day 56 vs. day 70 | -1.094 | -1.670 to -0.5181 | *** | <.001 | 10 | 10 |

**Figure 2G**

Kruskal-Wallis test

| | |
|---|---|
| P value | <.001 |
| Exact or approximate P value? | Approximate |
| P value summary | *** |
| Do the medians vary signif. (P < 0.05)? | Yes |
| Number of groups | 4 |
| Kruskal-Wallis statistic | 22.32 |

| | |
|---|---|
| Number of families | 1 |
| Number of comparisons per family | 3 |
| Alpha | 0.05 |

| Dunn's multiple comparisons test | Mean rank di | Significant? | Summary | Adjusted P Value | n1 | n2 |
|---|---|---|---|---|---|---|
| veh + veh vs. veh + subCGRP | 17.1 | Yes | ** | 0.003 | 10 | 9 |
| veh + subCGRP vs. PZN + subCGRP | -18.45 | Yes | ** | 0.001 | 9 | 10 |
| PZN + veh vs. PZN + subCGRP | 4.5 | No | ns | >.999 | 10 | 10 |

### Figure 3C

Kruskal-Wallis test

| | |
|---|---|
| Number of families | 1 |
| Number of comparisons per family | 6 |
| Alpha | 0.05 |

| | |
|---|---|
| P value | <.001 |
| Exact or approximate P value? | Approximate |
| P value summary | *** |
| Do the medians vary signif. (P < 0.05)? | Yes |
| Number of groups | 4 |
| Kruskal-Wallis statistic | 28.33 |

| Dunn's multiple comparisons test | Mean rank diff. | Significant? | Summary | Adjusted P Value | n1 | n2 |
|---|---|---|---|---|---|---|
| veh +veh vs. veh + subCGRP | 25.04 | Yes | *** | <.001 | 12 | 12 |
| veh +veh vs. PZN + veh | 14.71 | No | ns | 0.058 | 12 | 12 |
| veh +veh vs. PZN + subCGRP | 26.92 | Yes | *** | <.001 | 12 | 12 |
| veh + subCGRP vs. PZN + veh | -10.33 | No | ns | 0.413 | 12 | 12 |
| veh + subCGRP vs. PZN + subCGRF | 1.875 | No | ns | >.999 | 12 | 12 |
| PZN + veh vs. PZN + subCGRP | 12.21 | No | ns | 0.189 | 12 | 12 |

### Figure 3D

Kruskal-Wallis test

| | |
|---|---|
| Number of families | 1 |
| Number of comparisons per family | 6 |
| Alpha | 0.05 |

| | |
|---|---|
| P value | 0.004 |
| Exact or approximate P value? | Approximate |
| P value summary | ** |
| Do the medians vary signif. (P < 0.05)? | Yes |
| Number of groups | 4 |
| Kruskal-Wallis statistic | 13.1 |

| Dunn's multiple comparisons test | Mean rank diff. | Significant? | Summary | Adjusted P Value | n1 | n2 |
|---|---|---|---|---|---|---|
| veh +veh vs. veh + subCGRP | 11.42 | Yes | * | 0.03 | 6 | 6 |
| veh +veh vs. PZN + veh | 10.58 | No | ns | 0.055 | 6 | 6 |
| veh +veh vs. PZN + subCGRP | 13.33 | Yes | ** | 0.006 | 6 | 6 |
| veh + subCGRP vs. PZN + veh | -0.8333 | No | ns | >.999 | 6 | 6 |
| veh + subCGRP vs. PZN + subCGRF | 1.917 | No | ns | >.999 | 6 | 6 |
| PZN + veh vs. PZN + subCGRP | 2.75 | No | ns | >.999 | 6 | 6 |

### Figure 3E

Kruskal-Wallis test

| | |
|---|---|
| Number of families | 1 |
| Number of comparisons per family | 6 |
| Alpha | 0.05 |

| | |
|---|---|
| P value | 0.001 |
| Exact or approximate P value? | Approximate |
| P value summary | ** |
| Do the medians vary signif. (P < 0.05)? | Yes |
| Number of groups | 4 |
| Kruskal-Wallis statistic | 15.56 |

| Dunn's multiple comparisons test | Mean rank diff. | Significant? | Summary | Adjusted P Value | n1 | n2 |
|---|---|---|---|---|---|---|
| veh +veh vs. veh + subCGRP | 12.67 | Yes | * | 0.01 | 6 | 6 |
| veh +veh vs. PZN + veh | 4.75 | No | ns | >.999 | 6 | 6 |
| veh +veh vs. PZN + subCGRP | 13.58 | Yes | ** | 0.005 | 6 | 6 |
| veh + subCGRP vs. PZN + veh | -7.917 | No | ns | 0.302 | 6 | 6 |
| veh + subCGRP vs. PZN + subCGRF | 0.9167 | No | ns | >.999 | 6 | 6 |
| PZN + veh vs. PZN + subCGRP | 8.833 | No | ns | 0.174 | 6 | 6 |

### Figure 3G

Kruskal-Wallis test

| | |
|---|---|
| Number of families | 1 |
| Number of comparisons per family | 6 |
| Alpha | 0.05 |

| | |
|---|---|
| P value | <.001 |
| Exact or approximate P value? | Approximate |
| P value summary | *** |
| Do the medians vary signif. (P < 0.05)? | Yes |
| Number of groups | 4 |
| Kruskal-Wallis statistic | 27.97 |

| Dunn's multiple comparisons test | Mean rank diff. | Significant? | Summary | Adjusted P Value | n1 | n2 |
|---|---|---|---|---|---|---|
| veh +veh vs. veh + subCGRP | 21.46 | Yes | *** | <.001 | 12 | 12 |
| veh +veh vs. PZN + veh | 24.29 | Yes | *** | <.001 | 12 | 12 |
| veh +veh vs. PZN + subCGRP | 26.25 | Yes | *** | <.001 | 12 | 12 |
| veh + subCGRP vs. PZN + veh | 2.833 | No | ns | >.999 | 12 | 12 |
| veh + subCGRP vs. PZN + subCGRF | 4.792 | No | ns | >.999 | 12 | 12 |
| PZN + veh vs. PZN + subCGRP | 1.958 | No | ns | >.999 | 12 | 12 |

### Figure 3H

Kruskal-Wallis test

| | |
|---|---|
| Number of families | 1 |
| Number of comparisons per family | 6 |
| Alpha | 0.05 |

| | |
|---|---|
| P value | 0.003 |
| Exact or approximate P value? | Approximate |
| P value summary | ** |
| Do the medians vary signif. (P < 0.05)? | Yes |
| Number of groups | 4 |
| Kruskal-Wallis statistic | 13.62 |

| Dunn's multiple comparisons test | Mean rank diff. | Significant? | Summary | Adjusted P Value | n1 | n2 |
|---|---|---|---|---|---|---|

| | | | | | | |
|---|---|---|---|---|---|---|
| veh +veh vs. veh + subCGRP | 13.17 | Yes | ** | 0.006 | 6 | 6 |
| veh +veh vs. PZN + veh | 11.33 | Yes | * | 0.029 | 6 | 6 |
| veh +veh vs. PZN + subCGRP | 11.5 | Yes | * | 0.025 | 6 | 6 |
| veh + subCGRP vs. PZN + veh | -1.833 | No | ns | >.999 | 6 | 6 |
| veh + subCGRP vs. PZN + subCGRP | -1.667 | No | ns | >.999 | 6 | 6 |
| PZN + veh vs. PZN + subCGRP | 0.1667 | No | ns | >.999 | 6 | 6 |

### Figure 3I

Kruskal-Wallis test

| | |
|---|---|
| P value | 0.001 |
| Exact or approximate P value? | Approximate |
| P value summary | ** |
| Do the medians vary signif. (P < 0.05)? | Yes |
| Number of groups | 4 |
| Kruskal-Wallis statistic | 15.98 |

| | |
|---|---|
| Number of families | 1 |
| Number of comparisons per family | 6 |
| Alpha | 0.05 |

| Dunn's multiple comparisons test | Mean rank diff. | Significant? | Summary | Adjusted P Value | n1 | n2 |
|---|---|---|---|---|---|---|
| veh +veh vs. veh + subCGRP | 8.417 | No | ns | 0.219 | 6 | 6 |
| veh +veh vs. PZN + veh | 12.75 | Yes | ** | 0.009 | 6 | 6 |
| veh +veh vs. PZN + subCGRP | 14.83 | Yes | ** | 0.001 | 6 | 6 |
| veh + subCGRP vs. PZN + veh | 4.333 | No | ns | >.999 | 6 | 6 |
| veh + subCGRP vs. PZN + subCGRP | 6.417 | No | ns | 0.665 | 6 | 6 |
| PZN + veh vs. PZN + subCGRP | 2.083 | No | ns | >.999 | 6 | 6 |

### Figure 4B

Mixed-effect analysis

| | |
|---|---|
| Number of families | 7 |
| Number of comparisons per row family | 6 |
| Number of comparisons per column family | 3 |
| Alpha | 0.05 |

| Fixed effects (type III) | P value | P value summary | Statistically significant (P < 0.05)? | F (DFn, DFd) | Geisser-Greenhouse's epsilon |
|---|---|---|---|---|---|
| Time | <.001 | *** | Yes | F (1.608, 146.3) = 369.5 | 0.8039 |
| Treatment Factor | <.001 | *** | Yes | F (3, 92) = 29.22 | |
| Time x Treatment Factor | <.001 | *** | Yes | F (4.824, 146.3) = 26.69 | 0.8039 |

| Tukey's multiple comparisons test | Mean diff. | 95.00% CI of diff. | Summary | Adjusted P Value | n1 | n2 |
|---|---|---|---|---|---|---|
| B | | | | | | |
| PPL + veh vs. veh + veh | -0.0795 | -0.3985 to 0.2395 | ns | 0.91 | 24 | 24 |
| PPL + subCGRP vs. veh + veh | -0.01334 | -0.3147 to 0.2880 | ns | >.999 | 24 | 24 |
| veh + veh vs. veh + subCGRP | 0.1231 | -0.2308 to 0.4769 | ns | 0.789 | 24 | 24 |
| Day 1 | | | | | | |
| PPL + veh vs. veh + veh | -0.01484 | -0.05169 to 0.02200 | ns | 0.703 | 23 | 24 |
| PPL + subCGRP vs. veh + veh | -0.01788 | -0.05422 to 0.01846 | ns | 0.555 | 24 | 24 |
| veh + veh vs. veh + subCGRP | 0.007379 | -0.02915 to 0.04391 | ns | 0.948 | 24 | 24 |
| Day 14 | | | | | | |
| PPL + veh vs. veh + veh | -0.8046 | -1.255 to -0.3545 | *** | <.001 | 23 | 24 |
| PPL + subCGRP vs. veh + veh | -1.41 | -1.633 to -1.187 | *** | <.001 | 24 | 24 |
| veh + veh vs. veh + subCGRP | 1.365 | 1.090 to 1.641 | *** | <.001 | 24 | 24 |
| veh + veh | | | | | | |
| B vs. day 1 | 1.419 | 1.221 to 1.616 | *** | <.001 | 24 | 24 |
| Day 1 vs. day 14 | -1.446 | -1.642 to -1.250 | *** | <.001 | 24 | 24 |
| veh + subCGRP | | | | | | |
| B vs. day 1 | 1.303 | 1.041 to 1.565 | *** | <.001 | 24 | 24 |
| Day 1 vs. day 14 | -0.08824 | -0.2571 to 0.08061 | ns | 0.405 | 24 | 24 |
| PPL + veh | | | | | | |
| B vs. D1 | 1.344 | 1.116 to 1.573 | *** | <.001 | 23 | 23 |
| D1 vs. W2 | -0.6161 | -0.9876 to -0.2446 | ** | 0.001 | 22 | 22 |
| PPL + subCGRP | | | | | | |
| B vs. day 1 | 1.423 | 1.226 to 1.621 | *** | <.001 | 24 | 24 |
| Day 1 vs. day 14 | -0.05421 | -0.09155 to -0.01686 | ** | 0.004 | 24 | 24 |

### Figure 4C

Kruskal-Wallis test

| | |
|---|---|
| Number of families | 1 |
| Number of comparisons per family | 6 |
| Alpha | 0.05 |

| | |
|---|---|
| P value | <.001 |
| Exact or approximate P value? | Approximate |
| P value summary | *** |
| Do the medians vary signif. (P < 0.05)? | Yes |
| Number of groups | 4 |
| Kruskal-Wallis statistic | 50.88 |

| Dunn's multiple comparisons test | Mean rank di | Significant? | Summary | Adjusted P Value | n1 | n2 |
|---|---|---|---|---|---|---|
| veh + veh vs. veh + subCGRP | 48.9 | Yes | *** | <.001 | 24 | 24 |
| veh + veh vs. PPL+ veh | 21.33 | Yes | * | 0.042 | 24 | 23 |
| veh + veh vs. PPL + subCGRP | 45.29 | Yes | *** | <.001 | 24 | 24 |
| veh + subCGRP vs. PPL+ veh | -27.57 | Yes | ** | 0.003 | 24 | 23 |
| veh + subCGRP vs. PPL + subCGRP | -3.604 | No | ns | >.999 | 24 | 24 |
| PPL+ veh vs. PPL + subCGRP | 23.96 | Yes | * | 0.015 | 23 | 24 |

### Figure 4D

Kruskal-Wallis test

| | |
|---|---|
| Number of families | 1 |
| Number of comparisons per family | 6 |
| Alpha | 0.05 |

| | |
|---|---|
| P value | <.001 |
| Exact or approximate P value? | Approximate |
| P value summary | *** |
| Do the medians vary signif. (P < 0.05)? | Yes |
| Number of groups | 4 |
| Kruskal-Wallis statistic | 33.8 |

| Dunn's multiple comparisons test | Mean rank di | Significant? | Summary | Adjusted P Value | n1 | n2 |
|---|---|---|---|---|---|---|
| veh + veh vs. veh + subCGRP | 23.63 | Yes | *** | <.001 | 12 | 12 |
| veh + veh vs. PPL + veh | 5.417 | No | ns | >.999 | 12 | 12 |
| veh + veh vs. PPL + subCGRP | 26.29 | Yes | *** | <.001 | 12 | 12 |
| veh + subCGRP vs. PPL + veh | -18.21 | Yes | ** | 0.006 | 12 | 12 |
| veh + subCGRP vs. PPL + subCGRP | 2.667 | No | ns | >.999 | 12 | 12 |
| PPL + veh vs. PPL + subCGRP | 20.88 | Yes | *** | <.001 | 12 | 12 |

### Figure 5B

Mixed effect analysis

| | |
|---|---|
| Number of families | 12 |
| Number of comparisons per row family | 6 |
| Number of comparisons per column family | 28 |
| Alpha | 0.05 |

| Fixed effects (type III) | P value | P value summary | Statistically significant (P < 0.05)? | F (DFn, DFd) | Geisser-Greenhouse's epsilon |
|---|---|---|---|---|---|
| Time | <.001 | *** | Yes | F (5.358, 450.8) = 34.81 | 0.7654 |
| Treatment Factor | <.001 | *** | Yes | F (3, 85) = 25.86 | |
| Time x Treatment Factor | <.001 | *** | Yes | F (16.07, 450.8) = 16.44 | 0.7654 |

| Tukey's multiple comparisons test | Mean diff. | 95.00% CI of diff. | Summary | Adjusted P Value | n1 | n2 |
|---|---|---|---|---|---|---|
| B | | | | | | |
| Sham + subCGRP vs. Sham + full CGRP | -0.152 | -0.5362 to 0.2321 | ns | 0.708 | 21 | 21 |
| TBI + veh vs. TBI + subCGRP | 0.006122 | -0.4721 to 0.4843 | ns | >.999 | 22 | 23 |
| Day1 | | | | | | |
| Sham + subCGRP vs. Sham + full CGRP | 0.02571 | -0.4408 to 0.4923 | ns | 0.999 | 21 | 22 |
| TBI + veh vs. TBI + subCGRP | 0.2258 | -0.09566 to 0.5473 | ns | 0.237 | 22 | 23 |
| Day 14 | | | | | | |
| Sham + subCGRP vs. Sham + full CGRP | 1.175 | 0.8738 to 1.475 | *** | <.001 | 21 | 22 |
| TBI + veh vs. TBI + subCGRP | 1.076 | 0.6993 to 1.452 | *** | <.001 | 21 | 23 |
| Day 28 | | | | | | |
| Sham + subCGRP vs. Sham + full CGRP | 0.8562 | 0.4521 to 1.260 | *** | <.001 | 21 | 22 |
| TBI + veh vs. TBI + subCGRP | 0.5124 | 0.08346 to 0.9414 | * | 0.014 | 22 | 23 |
| Day 42 | | | | | | |
| Sham + subCGRP vs. Sham + full CGRP | 0.8213 | 0.4014 to 1.241 | *** | <.001 | 22 | 22 |
| TBI + veh vs. TBI + subCGRP | 0.5224 | 0.09967 to 0.9451 | * | 0.01 | 22 | 23 |
| Day 56 | | | | | | |
| Sham + subCGRP vs. Sham + full CGRP | 0.4114 | 0.01329 to 0.8095 | * | 0.041 | 22 | 22 |
| TBI + veh vs. TBI + subCGRP | 0.08755 | -0.2901 to 0.4652 | ns | 0.925 | 22 | 23 |
| Day 70 | | | | | | |
| Sham + subCGRP vs. Sham + full CGRP | 1.293 | 0.9674 to 1.618 | *** | <.001 | 22 | 22 |
| TBI + veh vs. TBI + subCGRP | 1.049 | 0.6324 to 1.465 | *** | <.001 | 22 | 23 |
| Day 84 | | | | | | |
| Sham + subCGRP vs. Sham + full CGRP | 0.1856 | -0.2248 to 0.5959 | ns | 0.623 | 22 | 22 |
| TBI + veh vs. TBI + subCGRP | 0.1 | -0.2556 to 0.4556 | ns | 0.875 | 22 | 23 |
| TBI + veh | | | | | | |
| B vs. day 1 | 0.945 | 0.3471 to 1.543 | *** | <.001 | 22 | 22 |
| Day 1 vs. day 14 | -1.025 | -1.609 to -0.4413 | *** | <.001 | 21 | 21 |
| Day 14 vs. day 28 | -0.1385 | -0.4865 to 0.2094 | ns | 0.872 | 21 | 21 |
| Day 28 vs. day 42 | -0.02805 | -0.3838 to 0.3277 | ns | >.999 | 22 | 22 |
| Day 42 vs. day 56 | 0.09177 | -0.08089 to 0.2644 | ns | 0.637 | 22 | 22 |
| Day 56 vs. day 70 | 0.09394 | -0.3693 to 0.5572 | ns | 0.997 | 22 | 22 |
| Day 70 vs. day 84 | -0.1689 | -0.5497 to 0.2119 | ns | 0.805 | 22 | 22 |
| TBI + subCGRP | | | | | | |
| B vs. day 1 | 1.165 | 0.7444 to 1.585 | *** | <.001 | 23 | 23 |
| Day 1 vs. day 14 | -0.1835 | -0.4712 to 0.1041 | ns | 0.427 | 23 | 23 |
| Day 14 vs. day 28 | -0.7084 | -1.159 to -0.2574 | *** | <.001 | 23 | 23 |
| Day 28 vs. day 42 | -0.01808 | -0.4433 to 0.4072 | ns | >.999 | 23 | 23 |
| Day 42 vs. day 56 | -0.3431 | -0.7529 to 0.06683 | ns | 0.147 | 23 | 23 |
| Day 56 vs. day 70 | 1.055 | 0.6064 to 1.503 | *** | <.001 | 23 | 23 |
| Day 70 vs. day 84 | -1.117 | -1.582 to -0.6525 | *** | <.001 | 23 | 23 |
| Sham + subCGRP | | | | | | |
| B vs. day 1 | -0.0719 | -0.2462 to 0.1024 | ns | 0.847 | 20 | 20 |
| Day 1 vs. day 14 | -0.08812 | -0.5632 to 0.3870 | ns | 0.998 | 21 | 21 |
| Day 14 vs. day 28 | -0.1207 | -0.4106 to 0.1691 | ns | 0.845 | 21 | 21 |
| Day 28 vs. day 42 | 0.00409 | -0.08512 to 0.09330 | ns | >.999 | 21 | 21 |
| Day 42 vs. day 56 | -0.01822 | -0.1875 to 0.1511 | ns | >.999 | 22 | 22 |
| Day 56 vs. day 70 | 0.05838 | -0.05083 to 0.1676 | ns | 0.631 | 22 | 22 |
| Day 70 vs. day 84 | 0.03205 | -0.3108 to 0.3749 | ns | >.999 | 22 | 22 |
| Sham + full CGRP | | | | | | |
| B vs. day 1 | 0.1931 | -0.1624 to 0.5487 | ns | 0.608 | 21 | 21 |
| Day 1 vs. day 14 | 1.061 | 0.6267 to 1.495 | *** | <.001 | 22 | 22 |
| Day 14 vs. day 28 | -0.4392 | -0.9206 to 0.04228 | ns | 0.091 | 22 | 22 |
| Day 28 vs. day 42 | -0.03548 | -0.2470 to 0.1760 | ns | >.999 | 22 | 22 |
| Day 42 vs. day 56 | -0.4281 | -0.8688 to 0.01252 | ns | 0.061 | 22 | 22 |
| Day 56 vs. day 70 | 0.9395 | 0.4615 to 1.418 | *** | <.001 | 22 | 22 |
| Day 70 vs. day 84 | -1.075 | -1.522 to -0.6283 | *** | <.001 | 22 | 22 |

### Figure 5C

Kruskal-Wallis test

| | |
|---|---|
| Number of families | 1 |
| Number of comparisons per family | 3 |
| Alpha | 0.05 |

| | |
|---|---|
| P value | <.001 |
| Exact or approximate P value? | Approximate |
| P value summary | *** |
| Do the medians vary signif. (P < 0.05)? | Yes |
| Number of groups | 4 |
| Kruskal-Wallis statistic | 55.63 |

| Dunn's multiple comparisons test | Mean rank diff. | Significant? | Summary | Adjusted P Value | n1 | n2 |
|---|---|---|---|---|---|---|
| Sham + subCGRP vs. Sham + full CGRP | 39.87 | Yes | *** | <.001 | 21 | 22 |
| TBI + veh vs. TBI + subCGRP | 38.35 | Yes | *** | <.001 | 20 | 23 |
| Sham + full CGRP vs. TBI + subCGRP | -1.007 | No | ns | >.999 | 22 | 23 |

### Figure 5D

Kruskal-Wallis test

| | |
|---|---|
| Number of families | 1 |
| Number of comparisons per family | 3 |
| Alpha | 0.05 |

| | |
|---|---|
| P value | 0.096 |
| Exact or approximate P value? | Approximate |
| P value summary | ns |
| Do the medians vary signif. (P < 0.05)? | No |
| Number of groups | 4 |
| Kruskal-Wallis statistic | 6.355 |

| Dunn's multiple comparisons test | Mean rank diff. | Significant? | Summary | Adjusted P Value | n1 | n2 |
|---|---|---|---|---|---|---|
| Sham + subCGRP vs. Sham + full CGRP | 16.09 | Yes | * | 0.036 | 22 | 22 |
| TBI + veh vs. TBI + subCGRP | -0.6877 | No | ns | >.999 | 22 | 23 |
| Sham + full CGRP vs. TBI + subCGRP | -7.438 | No | ns | 0.722 | 22 | 23 |

### Figure 5E

Mixed-effect analysis

| | |
|---|---|
| Number of families | 12 |
| Number of comparisons per row family | 6 |
| Number of comparisons per column family | 28 |
| Alpha | 0.05 |

| Fixed effects (type III) | P value | P value summary | Statistically significant (P < 0.05)? | F (DFn, DFd) | Geisser-Greenhouse's epsilon |
|---|---|---|---|---|---|
| Time | <.001 | *** | Yes | F (3.726, 138.4) = 16.55 | 0.5323 |
| Treatment Factor | <.001 | *** | Yes | F (3, 38) = 6.758 | |
| Time x Treatment Factor | <.001 | *** | Yes | F (11.18, 138.4) = 4.984 | 0.5323 |

| Tukey's multiple comparisons test | Mean diff. | 95.00% CI of diff. | Summary | Adjusted P Value | n1 | n2 |
|---|---|---|---|---|---|---|
| B | | | | | | |
| Sham + subCGRP vs. Sham + full CGRP | -0.1461 | -0.6633 to 0.3711 | ns | 0.837 | 9 | 9 |
| TBI + veh vs. TBI + subCGRP | 0.07779 | -0.5701 to 0.7257 | ns | 0.987 | 11 | 11 |
| Day 1 | | | | | | |
| Sham + subCGRP vs. Sham + full CGRP | -0.1246 | -0.8584 to 0.6093 | ns | 0.959 | 9 | 10 |
| TBI + veh vs. TBI + subCGRP | -0.1687 | -1.033 to 0.6953 | ns | 0.946 | 11 | 11 |
| Day 14 | | | | | | |
| Sham + subCGRP vs. Sham + full CGRP | 1.108 | 0.5425 to 1.673 | *** | <.001 | 9 | 10 |
| TBI + veh vs. TBI + subCGRP | 0.7948 | 0.04291 to 1.547 | * | 0.036 | 10 | 11 |
| Day 28 | | | | | | |
| Sham + subCGRP vs. Sham + full CGRP | 0.4995 | -0.2622 to 1.261 | ns | 0.274 | 9 | 10 |
| TBI + veh vs. TBI + subCGRP | 0.7609 | 0.03081 to 1.491 | * | 0.039 | 11 | 11 |

| | | | | | | |
|---|---|---|---|---|---|---|
| Day 42 | | | | | | |
| Sham + subCGRP vs. Sham + full CGRP | 0.3857 | -0.3467 to 1.118 | ns | 0.451 | 10 | 10 |
| TBI + veh vs. TBI + subCGRP | 0.3667 | -0.1187 to 0.8521 | ns | 0.16 | 11 | 11 |
| Day 56 | | | | | | |
| Sham + subCGRP vs. Sham + full CGRP | 0.2039 | -0.3400 to 0.7479 | ns | 0.7 | 10 | 10 |
| TBI + veh vs. TBI + subCGRP | -0.0007818 | -0.3081 to 0.3065 | ns | >.999 | 11 | 11 |
| Day 70 | | | | | | |
| Sham + subCGRP vs. Sham + full CGRP | 1.206 | 0.6239 to 1.788 | *** | <.001 | 10 | 10 |
| TBI + veh vs. TBI + subCGRP | 0.8041 | 0.01030 to 1.598 | * | 0.046 | 11 | 11 |
| Day 84 | | | | | | |
| Sham + subCGRP vs. Sham + full CGRP | 0.02097 | -0.4452 to 0.4872 | ns | >.999 | 10 | 10 |
| TBI + veh vs. TBI + subCGRP | 0.11 | -0.2105 to 0.4304 | ns | 0.726 | 11 | 11 |
| TBI + veh | | | | | | |
| B vs. day 1 | 0.8563 | -0.2553 to 1.968 | ns | 0.174 | 11 | 11 |
| Day 1 vs. day 14 | -0.658 | -1.872 to 0.5557 | ns | 0.486 | 10 | 10 |
| Day 14 vs. day 28 | -0.2436 | -0.7638 to 0.2767 | ns | 0.636 | 10 | 10 |
| Day 28 vs. day 42 | -0.1414 | -0.6716 to 0.3889 | ns | 0.964 | 11 | 11 |
| Day 42 vs. day 56 | 0.08052 | -0.2215 to 0.3825 | ns | 0.964 | 11 | 11 |
| Day 56 vs. day 70 | 0.2491 | -0.5208 to 1.019 | ns | 0.91 | 11 | 11 |
| Day 70 vs. day 84 | -0.3297 | -0.9830 to 0.3237 | ns | 0.583 | 11 | 11 |
| TBI + subCGRP | | | | | | |
| B vs. day 1 | 0.6098 | -0.2017 to 1.421 | ns | 0.192 | 11 | 11 |
| Day 1 vs. day 14 | 0.2662 | -1.061 to 1.593 | ns | 0.992 | 11 | 11 |
| Day 14 vs. day 28 | -0.2917 | -0.8645 to 0.2812 | ns | 0.573 | 11 | 11 |
| Day 28 vs. day 42 | -0.5355 | -1.542 to 0.4710 | ns | 0.527 | 11 | 11 |
| Day 42 vs. day 56 | -0.287 | -0.8597 to 0.2858 | ns | 0.59 | 11 | 11 |
| Day 56 vs. day 70 | 1.054 | 0.2157 to 1.892 | * | 0.012 | 11 | 11 |
| Day 70 vs. day 84 | -1.024 | -1.879 to -0.1687 | * | 0.017 | 11 | 11 |
| Sham + subCGRP | | | | | | |
| B vs. day 1 | -0.0803 | -0.5878 to 0.4272 | ns | 0.996 | 8 | 8 |
| Day 1 vs. day 14 | -0.001422 | -1.042 to 1.039 | ns | >.999 | 9 | 9 |
| Day 14 vs. day 28 | -0.1766 | -0.9047 to 0.5515 | ns | 0.969 | 9 | 9 |
| Day 28 vs. day 42 | 0 | | | | 9 | 9 |
| Day 42 vs. day 56 | -0.03755 | -0.2059 to 0.1308 | ns | 0.983 | 10 | 10 |
| Day 56 vs. day 70 | 0.07905 | -0.1267 to 0.2848 | ns | 0.803 | 10 | 10 |
| Day 70 vs. day 84 | -0.06114 | -0.7717 to 0.6495 | ns | >.999 | 10 | 10 |
| Sham + full CGRP | | | | | | |
| B vs. day 1 | 0.131 | -0.2467 to 0.5087 | ns | 0.847 | 9 | 9 |
| Day 1 vs. day 14 | 1.231 | 0.6814 to 1.781 | *** | <.001 | 10 | 10 |
| Day 14 vs. day 28 | -0.7849 | -1.583 to 0.01282 | ns | 0.054 | 10 | 10 |
| Day 28 vs. day 42 | -0.1285 | -0.6219 to 0.3650 | ns | 0.963 | 10 | 10 |
| Day 42 vs. day 56 | -0.2193 | -0.6855 to 0.2469 | ns | 0.631 | 10 | 10 |
| Day 56 vs. day 70 | 1.081 | 0.2781 to 1.884 | ** | 0.009 | 10 | 10 |
| Day 70 vs. day 84 | -1.246 | -1.947 to -0.5455 | ** | 0.001 | 10 | 10 |

### Figure 5F

Kruskal-Wallis test

| | |
|---|---|
| Number of families | 1 |
| Number of comparisons per family | 3 |
| Alpha | 0.05 |

| | |
|---|---|
| P value | <.001 |
| Exact or approximate P value? | Approximate |
| P value summary | *** |
| Do the medians vary signif. (P < 0.05)? | Yes |
| Number of groups | 4 |
| Kruskal-Wallis statistic | 21.38 |

| Dunn's multiple comparisons test | Mean rank diff. | Significant? | Summary | Adjusted P Value | n1 | n2 |
|---|---|---|---|---|---|---|
| Sham + subCGRP vs. Sham + full CGRP | 18.46 | Yes | *** | <.001 | 9 | 10 |
| Sham + full CGRP vs. TBI + subCGRP | -4.673 | No | ns | >.999 | 10 | 11 |
| TBI + veh vs. TBI + subCGRP | 14.06 | Yes | * | 0.015 | 9 | 11 |

### Figure 5G

Kruskal-Wallis test

| | |
|---|---|
| Number of families | 1 |
| Number of comparisons per family | 3 |
| Alpha | 0.05 |

| | |
|---|---|
| P value | 0.695 |
| Exact or approximate P value? | Approximate |
| P value summary | ns |
| Do the medians vary signif. (P < 0.05)? | No |
| Number of groups | 4 |
| Kruskal-Wallis statistic | 1.444 |

| Dunn's multiple comparisons test | Mean rank diff. | Significant? | Summary | Adjusted P Value | n1 | n2 |
|---|---|---|---|---|---|---|
| Sham + subCGRP vs. Sham + full CGRP | 2.35 | No | ns | >.999 | 10 | 10 |
| Sham + full CGRP vs. TBI + subCGRP | -2.859 | No | ns | >.999 | 10 | 11 |
| TBI + veh vs. TBI + subCGRP | 1.5 | No | ns | >.999 | 11 | 11 |

### Figure 5H

Two-way ANOVA

| | |
|---|---|
| Number of families | 12 |
| Number of comparisons per row family | 6 |
| Number of comparisons per column family | 28 |
| Alpha | 0.05 |

| Source of Variation | % of total vari: | P value | P value summary | Significant? | Geisser-Greenhouse's epsilon |
|---|---|---|---|---|---|
| Time x Treatment Factor | 23.34 | <.001 | *** | Yes | 0.7161 |
| Time | 10.41 | <.001 | *** | Yes | 0.7161 |
| Treatment Factor | 23.42 | <.001 | *** | Yes | |

| Tukey's multiple comparisons test | Mean diff. | 95.00% CI of diff. | Summary | Adjusted P Value | n1 | n2 |
|---|---|---|---|---|---|---|
| B | | | | | | |
| Sham + subCGRP vs. Sham + full CGRP | -0.1565 | -0.7653 to 0.4523 | ns | 0.884 | 12 | 12 |
| TBI + veh vs. TBI + subCGRP | -0.07078 | -0.8394 to 0.6978 | ns | 0.994 | 11 | 12 |
| Day 1 | | | | | | |
| Sham + subCGRP vs. Sham + full CGRP | 0.1496 | -0.5440 to 0.8432 | ns | 0.931 | 12 | 12 |
| TBI + veh vs. TBI + subCGRP | 0.01094 | -0.07789 to 0.09978 | ns | 0.986 | 11 | 12 |
| Day 14 | | | | | | |
| Sham + subCGRP vs. Sham + full CGRP | 1.222 | 0.8410 to 1.602 | *** | <.001 | 12 | 12 |
| TBI + veh vs. TBI + subCGRP | 1.369 | 1.070 to 1.667 | *** | <.001 | 11 | 12 |
| Day 28 | | | | | | |
| Sham + subCGRP vs. Sham + full CGRP | 1.149 | 0.7704 to 1.529 | *** | <.001 | 12 | 12 |
| TBI + veh vs. TBI + subCGRP | 0.8305 | 0.2601 to 1.401 | ** | 0.004 | 11 | 12 |
| Day 42 | | | | | | |
| Sham + subCGRP vs. Sham + full CGRP | 1.184 | 0.8092 to 1.559 | *** | <.001 | 12 | 12 |
| TBI + veh vs. TBI + subCGRP | 0.6557 | -0.03695 to 1.348 | ns | 0.068 | 11 | 12 |
| Day 56 | | | | | | |
| Sham + subCGRP vs. Sham + full CGRP | 0.5843 | -0.03643 to 1.205 | ns | 0.068 | 12 | 12 |
| TBI + veh vs. TBI + subCGRP | 0.1582 | -0.5145 to 0.8308 | ns | 0.912 | 11 | 12 |
| Day 70 | | | | | | |
| Sham + subCGRP vs. Sham + full CGRP | 1.365 | 0.9488 to 1.780 | *** | <.001 | 12 | 12 |
| TBI + veh vs. TBI + subCGRP | 1.275 | 0.8654 to 1.685 | *** | <.001 | 11 | 12 |
| Day 84 | | | | | | |
| Sham + subCGRP vs. Sham + full CGRP | 0.3227 | -0.3370 to 0.9824 | ns | 0.535 | 12 | 12 |
| TBI + veh vs. TBI + subCGRP | 0.08004 | -0.5743 to 0.7344 | ns | 0.986 | 11 | 12 |
| TBI + veh | | | | | | |
| B vs. day 1 | 1.034 | 0.2464 to 1.821 | ** | 0.009 | 11 | 11 |
| Day 1 vs. day 14 | -1.359 | -1.724 to -0.9942 | *** | <.001 | 11 | 11 |
| Day 14 vs. day 28 | -0.04306 | -0.6210 to 0.5349 | ns | >.999 | 11 | 11 |
| Day 28 vs. day 42 | 0.08528 | -0.5040 to 0.6745 | ns | 0.999 | 11 | 11 |
| Day 42 vs. day 56 | 0.103 | -0.1519 to 0.3580 | ns | 0.785 | 11 | 11 |
| Day 56 vs. day 70 | -0.06125 | -0.7443 to 0.6218 | ns | >.999 | 11 | 11 |
| Day 70 vs. day 84 | -0.008109 | -0.5199 to 0.5037 | ns | >.999 | 11 | 11 |
| TBI + subCGRP | | | | | | |
| B vs. day 1 | 1.115 | 0.4064 to 1.825 | ** | 0.002 | 12 | 12 |

| | | | | | | |
|---|---|---|---|---|---|---|
| Day 1 vs. day 14 | -0.001417 | -0.09145 to 0.08861 | ns | >.999 | 12 | 12 |
| Day 14 vs. day 28 | -0.5812 | -1.240 to 0.07737 | ns | 0.097 | 12 | 12 |
| Day 28 vs. day 42 | -0.08958 | -0.8795 to 0.7004 | ns | >.999 | 12 | 12 |
| Day 42 vs. day 56 | -0.3945 | -1.110 to 0.3211 | ns | 0.506 | 12 | 12 |
| Day 56 vs. day 70 | 1.056 | 0.4451 to 1.666 | *** | <.001 | 12 | 12 |
| Day 70 vs. day 84 | -1.203 | -1.836 to -0.5702 | *** | <.001 | 12 | 12 |
| Sham + subCGRP | | | | | | |
| B vs. day 1 | -0.06631 | -0.1906 to 0.05801 | ns | 0.542 | 12 | 12 |
| Day 1 vs. day 14 | -0.1531 | -0.7298 to 0.4235 | ns | 0.969 | 12 | 12 |
| Day 14 vs. day 28 | -0.07886 | -0.3307 to 0.1730 | ns | 0.93 | 12 | 12 |
| Day 28 vs. day 42 | 0.007158 | -0.1664 to 0.1807 | ns | >.999 | 12 | 12 |
| Day 42 vs. day 56 | -0.002108 | -0.3219 to 0.3177 | ns | >.999 | 12 | 12 |
| Day 56 vs. day 70 | 0.04116 | -0.1102 to 0.1926 | ns | 0.965 | 12 | 12 |
| Day 70 vs. day 84 | 0.1097 | -0.2938 to 0.5132 | ns | 0.965 | 12 | 12 |
| Sham + full CGRP | | | | | | |
| B vs. day 1 | 0.2397 | -0.3970 to 0.8765 | ns | 0.847 | 12 | 12 |
| Day 1 vs. day 14 | 0.9189 | 0.1775 to 1.660 | * | 0.013 | 12 | 12 |
| Day 14 vs. day 28 | -0.151 | -0.7453 to 0.4432 | ns | 0.975 | 12 | 12 |
| Day 28 vs. day 42 | 0.04202 | -0.09864 to 0.1827 | ns | 0.944 | 12 | 12 |
| Day 42 vs. day 56 | -0.6022 | -1.376 to 0.1721 | ns | 0.173 | 12 | 12 |
| Day 56 vs. day 70 | 0.8215 | 0.09980 to 1.543 | * | 0.023 | 12 | 12 |
| Day 70 vs. day 84 | -0.9322 | -1.623 to -0.2411 | ** | 0.007 | 12 | 12 |

### Figure 5I

Kruskal-Wallis test

| | | | | | | | | |
|---|---|---|---|---|---|---|---|---|
| | | | | | | | P value | <.001 |
| Number of families | 1 | | | | | | Exact or approximate P value? | Approximate |
| Number of comparisons per family | 3 | | | | | | P value summary | *** |
| Alpha | 0.05 | | | | | | Do the medians vary signif. (P < 0.05)? | Yes |
| | | | | | | | Number of groups | 4 |
| | | | | | | | Kruskal-Wallis statistic | 34.96 |
| Dunn's multiple comparisons test | Mean rank diff. | Significant? | Summary | Adjusted P Value | n1 | n2 | | |
| Sham + subCGRP vs. Sham + full CGRP | 22.17 | Yes | *** | <.001 | 12 | 12 | | |
| Sham + full CGRP vs. TBI + subCGRP | 2.083 | No | ns | >.999 | 12 | 12 | | |
| TBI + veh vs. TBI + subCGRP | 23.61 | Yes | *** | <.001 | 11 | 12 | | |

### Figure 5J

Kruskal-Wallis test

| | | | | | | | | |
|---|---|---|---|---|---|---|---|---|
| | | | | | | | P value | 0.045 |
| Number of families | 1 | | | | | | Exact or approximate P value? | Approximate |
| Number of comparisons per family | 3 | | | | | | P value summary | * |
| Alpha | 0.05 | | | | | | Do the medians vary signif. (P < 0.05)? | Yes |
| | | | | | | | Number of groups | 4 |
| | | | | | | | Kruskal-Wallis statistic | 8.03 |
| Dunn's multiple comparisons test | Mean rank diff. | Significant? | Summary | Adjusted P Value | n1 | n2 | | |
| Sham + subCGRP vs. Sham + full CGRP | 13.33 | Yes | * | 0.024 | 12 | 12 | | |
| Sham + full CGRP vs. TBI + subCGRP | -4.333 | No | ns | >.999 | 12 | 12 | | |
| TBI + veh vs. TBI + subCGRP | -2.064 | No | ns | >.999 | 11 | 12 | | |

**Figure 6D**
Ordinary One-Way ANOVA

| | | | | | | | |
|---|---|---|---|---|---|---|---|
| | | | | | | F | 3.152 |
| Number of comparisons per family | 2 | | | | | P value | 0.043 |
| Alpha | 0.05 | | | | | P value summary | * |
| | | | | | | Significant diff. among means (P < 0.05)? | Yes |
| | | | | | | R squared | 0.2744 |
| Holm-Sidak's multiple comparisons test | Mean diff. | Summary | Adjusted P Value | n1 | n2 | | |
| veh + veh vs. veh + full CGRP | 31.33 | * | 0.026 | 7 | 7 | | |
| veh + full CGRP vs. PZN + full CGRP | -37 | * | 0.016 | 7 | 8 | | |

**Figure 6E**
Ordinary One-Way ANOVA

| | | | | | | | |
|---|---|---|---|---|---|---|---|
| | | | | | | F | 1.278 |
| Number of families | 1 | | | | | P value | 0.304 |
| Number of comparisons per family | 2 | | | | | P value summary | ns |
| Alpha | 0.05 | | | | | Significant diff. among means (P < 0.05)? | No |
| | | | | | | R squared | 0.1329 |
| Holm-Sidak's multiple comparisons test | Mean diff. | Summary | Adjusted P Value | n1 | n2 | | |
| veh + veh vs. veh + full CGRP | 41.12 | ns | 0.15 | 7 | 7 | | |
| veh + full CGRP vs. PZN + full CGRP | -31.58 | ns | 0.157 | 7 | 8 | | |

**Figure 6F**
Ordinary One-Way ANOVA

| | | | | | | | |
|---|---|---|---|---|---|---|---|
| | | | | | | F | 1.693 |
| Number of families | 1 | | | | | P value | 0.194 |
| Number of comparisons per family | 2 | | | | | P value summary | ns |
| Alpha | 0.05 | | | | | Significant diff. among means (P < 0.05)? | No |
| | | | | | | R squared | 0.1689 |
| Holm-Sidak's multiple comparisons test | Mean diff. | Summary | Adjusted P Value | n1 | n2 | | |
| veh + veh vs. veh + full CGRP | 32.77 | ns | 0.181 | 7 | 7 | | |
| veh + full CGRP vs. PZN + full CGRP | -25.33 | ns | 0.181 | 7 | 8 | | |

**Figure 6G**
Ordinary One-Way ANOVA

| | | | | | | | |
|---|---|---|---|---|---|---|---|
| | | | | | | F | 0.8506 |
| Number of families | 1 | | | | | P value | 0.479 |
| Number of comparisons per family | 2 | | | | | P value summary | ns |
| Alpha | 0.05 | | | | | Significant diff. among means (P < 0.05)? | No |
| | | | | | | R squared | 0.09262 |
| Holm-Sidak's multiple comparisons test | Mean diff. | Summary | Adjusted P Value | n1 | n2 | | |
| veh + veh vs. veh + full CGRP | 15.62 | ns | 0.349 | 7 | 7 | | |
| veh + full CGRP vs. PZN + full CGRP | -24.28 | ns | 0.257 | 7 | 8 | | |

**Figure 6H**
Ordinary One-Way ANOVA

| | | | | | | | |
|---|---|---|---|---|---|---|---|
| | | | | | | F | 2.448 |
| Number of families | 1 | | | | | P value | 0.088 |
| Number of comparisons per family | 2 | | | | | P value summary | ns |
| Alpha | 0.05 | | | | | Significant diff. among means (P < 0.05)? | No |
| | | | | | | R squared | 0.2343 |
| Holm-Sidak's multiple comparisons test | Mean diff. | Summary | Adjusted P Value | n1 | n2 | | |
| veh + veh vs. veh + full CGRP | 61.63 | * | 0.026 | 6 | 7 | | |
| veh + full CGRP vs. PZN + full CGRP | -31.23 | ns | 0.292 | 7 | 8 | | |

**Figure 6I**
Kruskal-Wallis test

| | | | | | | | |
|---|---|---|---|---|---|---|---|
| | | | | | | P value | 0.311 |
| Number of families | 1 | | | | | Exact or approximate P value? | Approximate |
| Number of comparisons per family | 2 | | | | | P value summary | ns |
| Alpha | 0.05 | | | | | Do the medians vary signif. (P < 0.05)? | No |
| | | | | | | Number of groups | 4 |
| | | | | | | Kruskal-Wallis statistic | 3.575 |
| Dunn's multiple comparisons test | Mean rank diff. | Summary | Adjusted P Value | n1 | n2 | | |
| veh + veh vs. veh + full CGRP | 7.952 | ns | 0.165 | 6 | 7 | | |
| veh + full CGRP vs. PZN + full CGRP | -6.411 | ns | 0.264 | 7 | 8 | | |

### Figure 7A

Nonlin fit

| Best-fit values | AQP4 KO | WT |
|---|---|---|
| Bottom | 0.02016 | -0.03357 |
| Hillslope | Unstable | -3.241 |
| Top | 1.449 | 1.431 |
| EC50 | 0.004394 | 0.03356 |
| logEC50 | -2.357 | -1.474 |
| Span | 1.429 | 1.465 |
| Goodness of Fit | | |
| Degrees of Freedom | 108 | 104 |
| R squared | 0.841 | 0.6879 |
| Sum of Squares | 8.92 | 13.8 |
| Sy.x | 0.2874 | 0.3643 |

Wilcoxon matched-pairs signed rank test
AQP4 KO
CGRP 0.005 mg kg-1 vs veh

| | |
|---|---|
| P value | <.001 |
| Exact or approximate P value? | Exact |
| P value summary | *** |
| Significantly different (P < 0.05)? | Yes |
| One- or two-tailed P value? | Two-tailed |
| Sum of positive, negative ranks | 1.000 , -377.0 |
| Sum of signed ranks (W) | -376 |
| Number of pairs | 28 |
| Number of ties (ignored) | 1 |

Wilcoxon matched-pairs signed rank test
AQP4 KO
CGRP 0.01 mg kg-1 vs veh

| | |
|---|---|
| P value | <.001 |
| Exact or approximate P value? | Exact |
| P value summary | *** |
| Significantly different (P < 0.05)? | Yes |
| One- or two-tailed P value? | Two-tailed |
| Sum of positive, negative ranks | 0.000 , -406.0 |
| Sum of signed ranks (W) | -406 |
| Number of pairs | 28 |
| Number of ties (ignored) | 0 |

Wilcoxon matched-pairs signed rank test
WT
0.05 mg kg-1 vs veh

| | |
|---|---|
| P value | <.001 |
| Exact or approximate P value? | Exact |
| P value summary | *** |
| Significantly different (P < 0.05)? | Yes |
| One- or two-tailed P value? | Two-tailed |
| Sum of positive, negative ranks | 0.000 , -91.00 |
| Sum of signed ranks (W) | -91 |
| Number of pairs | 13 |
| Number of ties (ignored) | 0 |

Mann Whitney test
WT
0.1 mg kg-1 vs veh

| | |
|---|---|
| P value | <.001 |
| Exact or approximate P value? | Exact |
| P value summary | *** |
| Significantly different (P < 0.05)? | Yes |
| One- or two-tailed P value? | Two-tailed |
| Sum of ranks in column A,B | 462 , 66 |
| Mann-Whitney U | 0 |

### Figure 7C

Mixed-effect analysis

| | |
|---|---|
| Number of families | 8 |
| Number of comparisons per row family | 6 |
| Number of comparisons per column family | 6 |
| Alpha | 0.05 |

| Fixed effects (type III) | P value | P value summary | Statistically significant (P < 0.05)? | F (DFn, DFd) | Geisser-Greenhouse's epsilon |
|---|---|---|---|---|---|
| Time | <.001 | *** | Yes | F (2.353, 28.24) = 14.78 | 0.7845 |
| Treatment Factor | <.001 | *** | Yes | F (2.185, 26.22) = 47.03 | 0.7283 |
| Time x Treatment Factor | <.001 | *** | Yes | F (3.812, 38.97) = 13.24 | 0.4236 |

| Tukey's multiple comparisons test | Mean diff. | 95.00% CI of diff. | Summary | Adjusted P Value | n1 | n2 |
|---|---|---|---|---|---|---|
| B | | | | | | |
| WT + veh vs. WT + CGRP 0.1 mg kg-1 | 0 | | | | 11 | 11 |
| KO + veh vs. KO + CGRP 0.01 mg kg-1 | 0 | | | | 13 | 13 |
| Day 0 | | | | | | |
| WT + veh vs. WT + CGRP 0.1 mg kg-1 | 1.29 | 0.7785 to 1.801 | *** | <.001 | 11 | 11 |
| KO + veh vs. KO + CGRP 0.01 mg kg-1 | 0.961 | 0.4757 to 1.446 | *** | <.001 | 13 | 13 |
| Day 14 | | | | | | |
| WT + veh vs. WT + CGRP 0.1 mg kg-1 | 0.2848 | -0.3417 to 0.9113 | ns | 0.532 | 11 | 11 |
| KO + veh vs. KO + CGRP 0.01 mg kg-1 | 1.254 | 0.8791 to 1.628 | *** | <.001 | 13 | 13 |
| Day 42 | | | | | | |
| WT + veh vs. WT + CGRP 0.1 mg kg-1 | 0.07388 | -0.6333 to 0.7811 | ns | 0.988 | 11 | 11 |
| KO + veh vs. KO + CGRP 0.01 mg kg-1 | 1.36 | 0.9707 to 1.749 | *** | <.001 | 13 | 13 |
| WT + veh | | | | | | |
| B vs. day 0 | 0 | | | | 11 | 11 |
| Day 0 vs. day 14 | 0.03695 | -0.5531 to 0.6270 | ns | 0.997 | 11 | 11 |
| Day 14 vs. day 42 | -0.06886 | -0.5903 to 0.4526 | ns | 0.977 | 11 | 11 |
| WT + CGRP 0.1 mg kg-1 | | | | | | |
| B vs. day 0 | 1.29 | 0.7785 to 1.801 | *** | <.001 | 11 | 11 |
| Day 0 vs. day 14 | -0.9678 | -1.716 to -0.2195 | * | 0.012 | 11 | 11 |
| Day 14 vs. day 42 | -0.2798 | -1.081 to 0.5213 | ns | 0.715 | 11 | 11 |
| KO + veh | | | | | | |
| B vs. day 0 | 0 | | | | 13 | 13 |
| Day 0 vs. day 14 | 0.008692 | -0.4829 to 0.5002 | ns | >.999 | 13 | 13 |
| Day 14 vs. day 42 | -0.08839 | -0.5003 to 0.3235 | ns | 0.918 | 13 | 13 |
| KO + CGRP 0.01 mg kg-1 | | | | | | |
| B vs. day 0 | 0.961 | 0.4757 to 1.446 | *** | <.001 | 13 | 13 |
| Day 0 vs. day 14 | 0.3013 | -0.05613 to 0.6587 | ns | 0.11 | 13 | 13 |
| Day 14 vs. day 42 | 0.01765 | -0.06727 to 0.1026 | ns | 0.925 | 13 | 13 |

### Figure 7D

Kruskal-Wallis test

| | |
|---|---|
| Number of families | 1 |
| Number of comparisons per family | 3 |
| Alpha | 0.05 |

| | |
|---|---|
| P value | <.001 |
| Exact or approximate P value? | Approximate |
| P value summary | *** |
| Do the medians vary signif. (P < 0.05)? | Yes |
| Number of groups | 5 |
| Kruskal-Wallis statistic | 40.08 |

| Dunn's multiple comparisons test | Mean rank diff. | Significant? | Summary | Adjusted P Value | n1 | n2 |
|---|---|---|---|---|---|---|
| WT + veh vs. WT + CGRP 0.01 mg kg-1 | -1.045 | No | ns | >.999 | 11 | 11 |
| WT + veh vs. WT + CGRP 0.1 mg kg-1 | 32.82 | Yes | *** | <.001 | 11 | 11 |
| KO + veh vs. KO + CGRP 0.01 mg kg-1 | 21.54 | Yes | ** | 0.003 | 13 | 13 |

### Figure 7E

Kruskal-Wallis test

| | |
|---|---|
| Number of families | 1 |
| Number of comparisons per family | 2 |
| Alpha | 0.05 |

| | |
|---|---|
| P value | <.001 |
| Exact or approximate P value? | Approximate |
| P value summary | *** |
| Do the medians vary signif. (P < 0.05)? | Yes |
| Number of groups | 4 |
| Kruskal-Wallis statistic | 31.14 |

| Dunn's multiple comparisons test | Mean rank diff. | Significant? | Summary | Adjusted P Value | n1 | n2 |
|---|---|---|---|---|---|---|
| WT + veh vs. WT + CGRP 0.1 mg kg-1 | 1.773 | No | ns | >.999 | 11 | 11 |
| KO + veh vs. KO + CGRP 0.01 mg kg-1 | 24.19 | Yes | *** | <.001 | 13 | 13 |

### Figure 8B

Mixed-effect analysis

| | |
|---|---|
| Number of families | 8 |
| Number of comparisons per row family | 6 |
| Number of comparisons per column family | 6 |
| Alpha | 0.05 |

| Fixed effects (type III) | P value | P value summary | Statistically significant (P < 0.05)? | F (DFn, DFd) | Geisser-Greenhouse's epsilon |
|---|---|---|---|---|---|
| Treatment Factor | <.001 | *** | Yes | F (3, 52) = 49.04 | |
| Time | <.001 | *** | Yes | F (1.454, 51.38) = 43.40 | 0.4847 |
| Treatment Factor x Time | <.001 | *** | Yes | F (4.362, 51.38) = 15.27 | 0.4847 |

| Tukey's multiple comparisons test | Mean diff. | 95.00% CI of diff. | Summary | Adjusted P Value | n1 | n2 |
|---|---|---|---|---|---|---|
| B | | | | | | |
| WT + veh vs. WT + CGRP 0.001 mg kg-1 | 0 | | | | 8 | 8 |
| AQP4 KO + veh vs. AQP4 KO + CGRP 0.001 mg kg-1 | 0 | | | | 14 | 14 |
| Day 1 | | | | | | |
| WT + veh vs. WT + CGRP 0.001 mg kg-1 | 0 | | | | 8 | 8 |
| AQP4 KO + veh vs. AQP4 KO + CGRP 0.001 mg kg-1 | 0 | | | | 14 | 14 |
| Day 14 | | | | | | |
| WT + veh vs. WT + CGRP 0.001 mg kg-1 | -0.06169 | -0.4033 to 0.2799 | ns | 0.93 | 8 | 8 |
| AQP4 KO + veh vs. AQP4 KO + CGRP 0.001 mg kg-1 | 1.184 | 0.7479 to 1.620 | *** | <.001 | 14 | 14 |
| Day 56 | | | | | | |
| WT + veh vs. WT + CGRP 0.001 mg kg-1 | -0.1234 | -0.4957 to 0.2488 | ns | 0.677 | 7 | 7 |
| AQP4 KO + veh vs. AQP4 KO + CGRP 0.001 mg kg-1 | 1.421 | 1.094 to 1.748 | *** | <.001 | 14 | 14 |
| WT + veh | | | | | | |
| B vs. day 1 | 1.31 | 0.6949 to 1.924 | *** | <.001 | 8 | 8 |
| Day 1 vs. day 14 | -1.087 | -1.733 to -0.4410 | ** | 0.004 | 8 | 8 |
| Day 14 vs. day 56 | -0.179 | -0.9114 to 0.5535 | ns | 0.888 | 8 | 7 |
| WT + CGRP 0.001 mg kg-1 | | | | | | |
| B vs. day 1 | 1.31 | 0.6949 to 1.924 | *** | <.001 | 8 | 8 |
| Day 1 vs. day 14 | -1.149 | -1.870 to -0.4277 | ** | 0.005 | 8 | 8 |
| Day 14 vs. day 56 | -0.2407 | -1.004 to 0.5224 | ns | 0.782 | 8 | 7 |
| AQP4 KO + veh | | | | | | |
| B vs. day 1 | 1.284 | 0.8667 to 1.701 | *** | <.001 | 14 | 14 |
| Day 1 vs. day 14 | -1.172 | -1.614 to -0.7290 | *** | <.001 | 14 | 14 |
| Day 14 vs. day 56 | -0.2318 | -0.7477 to 0.2841 | ns | 0.609 | 14 | 14 |
| AQP4 KO + CGRP 0.001 mg kg-1 | | | | | | |
| B vs. day 1 | 1.284 | 0.8667 to 1.701 | *** | <.001 | 14 | 14 |
| Day 1 vs. day 14 | 0.0122 | -0.04192 to 0.06632 | ns | 0.917 | 14 | 14 |
| Day 14 vs. day 56 | 0.005564 | -0.01494 to 0.02606 | ns | 0.864 | 14 | 14 |

### Figure 8C

Kruskal-Wallis test

| | |
|---|---|
| Number of families | 1 |
| Number of comparisons per family | 2 |
| Alpha | 0.05 |

| | |
|---|---|
| P value | <.001 |
| Exact or approximate P value? | Approximate |
| P value summary | *** |
| Do the medians vary signif. (P < 0.05)? | Yes |
| Number of groups | 4 |
| Kruskal-Wallis statistic | 31.31 |

| Dunn's multiple comparisons test | Mean rank diff. | Significant? | Summary | Adjusted P Value | n1 | n2 |
|---|---|---|---|---|---|---|
| WT + veh vs. WT + CGRP 0.001 mg kg-1 | -2.357 | No | ns | >.999 | 7 | 7 |
| AQP4 KO + veh vs. AQP4 KO + CGRP 0.001 mg kg-1 | 21.82 | Yes | *** | <.001 | 14 | 14 |

### Figure 8E

Mixed-effect analysis

| | |
|---|---|
| Number of families | 9 |
| Number of comparisons per row family | 6 |
| Number of comparisons per column family | 10 |
| Alpha | 0.05 |

| Fixed effects (type III) | P value | P value summary | Statistically significant (P < 0.05)? | F (DFn, DFd) | Geisser-Greenhouse's epsilon |
|---|---|---|---|---|---|
| Time | <.001 | *** | Yes | F (2.789, 78.10) = 54.84 | 0.6973 |
| Treatment Factor | <.001 | *** | Yes | F (3, 44) = 13.47 | |
| Time x Treatment Factor | <.001 | *** | Yes | F (8.367, 78.10) = 7.257 | 0.6973 |

| Tukey's multiple comparisons test | Mean diff. | 95.00% CI of diff. | Summary | Adjusted P Value | n1 | n2 |
|---|---|---|---|---|---|---|
| B | | | | | | |
| veh + veh vs. veh + CGRP 0.01 mg kg-1 | 0 | -0.5910 to 0.5910 | ns | >.999 | 15 | 15 |
| veh + veh vs. PZN + veh | -0.1322 | -0.7312 to 0.4668 | ns | 0.926 | 15 | 9 |
| veh + veh vs. PZN + CGRP 0.01 mg kg-1 | -0.1322 | -0.7312 to 0.4668 | ns | 0.926 | 15 | 9 |
| Day 1 | | | | | | |
| veh + veh vs. veh + CGRP 0.01 mg kg-1 | 0 | -0.2789 to 0.2789 | ns | >.999 | 7 | 7 |
| veh + veh vs. PZN + veh | -0.01302 | -0.3430 to 0.3169 | ns | >.999 | 7 | 9 |
| veh + veh vs. PZN + CGRP 0.01 mg kg-1 | -0.01302 | -0.3430 to 0.3169 | ns | >.999 | 7 | 9 |
| Day 14 | | | | | | |
| veh + veh vs. veh + CGRP 0.01 mg kg-1 | 1.127 | 0.3399 to 1.915 | * | 0.01 | 7 | 7 |
| veh + veh vs. PZN + veh | 0.03023 | -0.8137 to 0.8742 | ns | >.999 | 7 | 9 |
| veh + veh vs. PZN + CGRP 0.01 mg kg-1 | 0.2475 | -0.6710 to 1.166 | ns | 0.859 | 7 | 9 |
| Day 28 | | | | | | |
| veh + veh vs. veh + CGRP 0.01 mg kg-1 | 1.036 | 0.1561 to 1.915 | * | 0.025 | 7 | 7 |
| veh + veh vs. PZN + veh | -0.4322 | -1.310 to 0.4453 | ns | 0.421 | 7 | 9 |
| veh + veh vs. PZN + CGRP 0.01 mg kg-1 | -0.3255 | -1.227 to 0.5762 | ns | 0.693 | 7 | 9 |
| Day 42 | | | | | | |
| veh + veh vs. veh + CGRP 0.01 mg kg-1 | 1.258 | 0.4952 to 2.020 | ** | 0.005 | 7 | 7 |
| veh + veh vs. PZN + veh | -0.2307 | -0.9952 to 0.5337 | ns | 0.777 | 7 | 9 |
| veh + veh vs. PZN + CGRP 0.01 mg kg-1 | -0.1429 | -0.9407 to 0.6550 | ns | 0.947 | 7 | 9 |
| veh + veh | | | | | | |
| B vs. day 1 | 1.118 | 0.2111 to 2.026 | * | 0.02 | 7 | 7 |
| Day 1 vs. day 14 | -1.065 | -1.848 to -0.2813 | * | 0.013 | 7 | 7 |
| Day 14 vs. day 28 | 0.07504 | -0.6447 to 0.7947 | ns | 0.994 | 7 | 7 |
| Day 28 vs. day 42 | -0.1885 | -1.127 to 0.7501 | ns | 0.935 | 7 | 7 |
| veh + CGRP 0.01 mg kg-1 | | | | | | |
| B vs. day 1 | 1.118 | 0.2111 to 2.026 | * | 0.02 | 7 | 7 |
| Day 1 vs. day 14 | 0.06247 | -0.1832 to 0.3081 | ns | 0.866 | 7 | 7 |
| Day 14 vs. day 28 | -0.01653 | -0.1155 to 0.08244 | ns | 0.965 | 7 | 7 |
| Day 28 vs. day 42 | 0.03329 | -0.1203 to 0.1868 | ns | 0.917 | 7 | 7 |
| PZN + veh | | | | | | |
| B vs. day 1 | 1.305 | 0.7442 to 1.865 | *** | <.001 | 9 | 9 |
| Day 1 vs. day 14 | -1.022 | -1.581 to -0.4624 | ** | 0.002 | 9 | 9 |
| Day 14 vs. day 28 | -0.3874 | -1.098 to 0.3232 | ns | 0.395 | 9 | 9 |
| Day 28 vs. day 42 | 0.01298 | -0.4380 to 0.4639 | ns | >.999 | 9 | 9 |
| PZN + CGRP 0.01 mg kg-1 | | | | | | |
| B vs. day 1 | 1.305 | 0.7442 to 1.865 | *** | <.001 | 9 | 9 |
| Day 1 vs. day 14 | -0.8043 | -1.494 to -0.1148 | * | 0.023 | 9 | 9 |
| Day 14 vs. day 28 | -0.4979 | -1.253 to 0.2568 | ns | 0.244 | 9 | 9 |
| Day 28 vs. day 42 | -0.005944 | -0.2135 to 0.2016 | ns | >.999 | 9 | 9 |

### Figure 8F

Kruskal-Wallis test

| | |
|---|---|
| Number of families | 1 |
| Number of comparisons per family | 3 |
| Alpha | 0.05 |

| | |
|---|---|
| P value | <.001 |
| Exact or approximate P value? | Approximate |
| P value summary | *** |
| Do the medians vary signif. (P < 0.05)? | Yes |
| Number of groups | 4 |
| Kruskal-Wallis statistic | 21.51 |

| Dunn's multiple comparisons test | Mean rank diff. | Significant? | Summary | Adjusted P Value | n1 | n2 |
|---|---|---|---|---|---|---|
| veh + veh vs. veh + CGRP 0.01 mg kg-1 | 14.71 | Yes | ** | 0.002 | 7 | 7 |
| veh + CGRP 0.01 mg kg-1 vs. PZN + CGRP 0.01 mg kg-1 | -15.83 | Yes | *** | <.001 | 7 | 9 |
| PZN + veh vs. PZN + CGRP 0.01 mg kg-1 | 1.333 | No | ns | >.999 | 9 | 9 |

### Figure 8G

Kruskal-Wallis test

| | |
|---|---|
| Number of families | 1 |
| Number of comparisons per family | 3 |
| Alpha | 0.05 |

| | |
|---|---|
| P value | <.001 |
| Exact or approximate P value? | Approximate |
| P value summary | *** |
| Do the medians vary signif. (P < 0.05)? | Yes |
| Number of groups | 4 |

| | |
|---|---|
| Kruskal-Wallis statistic | 42.88 |

| Dunn's multiple comparisons test | Mean rank diff. | Significant? | Summary | Adjusted P Value | n1 | n2 |
|---|---|---|---|---|---|---|
| veh + veh vs. veh + SNP 0.25 mg kg-1 | 34 | Yes | *** | <.001 | 19 | 19 |
| PZN + veh vs. PZN + SNP 0.25 mg kg-1 | 12.96 | No | ns | 0.198 | 22 | 21 |
| veh + SNP 0.25 mg kg-1 vs. PZN + SNP 0.25 mg kg-1 | -32.78 | Yes | *** | <.001 | 19 | 21 |

### Figure 8H

Mixed-effect analysis

| | |
|---|---|
| Number of families | 9 |
| Number of comparisons per row family | 6 |
| Number of comparisons per column family | 10 |
| Alpha | 0.05 |

| Fixed effects (type III) | P value | P value summary | Statistically significant (P < 0.05)? | F (DFn, DFd) | Geisser-Greenhouse's epsilon |
|---|---|---|---|---|---|
| Time | <.001 | *** | Yes | F (2.931, 167.0) = 35.56 | 0.7326 |
| Treatment Factor | <.001 | *** | Yes | F (3, 78) = 29.61 | |
| Time x Treatment Factor | <.001 | *** | Yes | F (8.792, 167.0) = 9.203 | 0.7326 |

| Tukey's multiple comparisons test | Mean diff. | 95.00% CI of diff. | Summary | Adjusted P Value | N1 | N2 |
|---|---|---|---|---|---|---|
| B | | | | | | |
| veh + veh vs. veh + CGRP 0.01 mg kg-1 | 0 | -0.4708 to 0.4708 | ns | >.999 | 29 | 29 |
| veh + veh vs. PZN + veh | 0.1524 | -0.4699 to 0.7746 | ns | 0.903 | 29 | 12 |
| veh + veh vs. PZN + CGRP 0.01 mg kg-1 | 0.1524 | -0.4699 to 0.7746 | ns | 0.903 | 29 | 12 |
| Day 1 | | | | | | |
| veh + veh vs. veh + CGRP 0.01 mg kg-1 | 0 | -0.2066 to 0.2066 | ns | >.999 | 29 | 29 |
| veh + veh vs. PZN + veh | 0.0424 | -0.1145 to 0.1993 | ns | 0.884 | 29 | 12 |
| veh + veh vs. PZN + CGRP 0.01 mg kg-1 | 0.0424 | -0.1145 to 0.1993 | ns | 0.884 | 29 | 12 |
| Day 14 | | | | | | |
| veh + veh vs. veh + CGRP 0.01 mg kg-1 | 0.979 | 0.4956 to 1.462 | *** | <.001 | 15 | 15 |
| veh + veh vs. PZN + veh | -0.1746 | -0.8117 to 0.4624 | ns | 0.874 | 15 | 12 |
| veh + veh vs. PZN + CGRP 0.01 mg kg-1 | 0.9681 | 0.4816 to 1.455 | *** | <.001 | 15 | 12 |
| Day 28 | | | | | | |
| veh + veh vs. veh + CGRP 0.01 mg kg-1 | 0.9743 | 0.5205 to 1.428 | *** | <.001 | 15 | 15 |
| veh + veh vs. PZN + veh | -0.1357 | -0.7508 to 0.4794 | ns | 0.929 | 15 | 12 |
| veh + veh vs. PZN + CGRP 0.01 mg kg-1 | 0.966 | 0.5120 to 1.420 | *** | <.001 | 15 | 12 |
| Day 42 | | | | | | |
| veh + veh vs. veh + CGRP 0.01 mg kg-1 | 0.9646 | 0.5066 to 1.423 | *** | <.001 | 15 | 15 |
| veh + veh vs. PZN + veh | -0.1938 | -0.8698 to 0.4822 | ns | 0.857 | 15 | 12 |
| veh + veh vs. PZN + CGRP 0.01 mg kg-1 | 0.9644 | 0.5064 to 1.422 | *** | <.001 | 15 | 12 |
| veh + veh | | | | | | |
| B vs. day 1 | 0.9332 | 0.5160 to 1.350 | *** | <.001 | 29 | 29 |
| Day 1 vs. day 14 | -0.8882 | -1.423 to -0.3535 | ** | 0.001 | 15 | 15 |
| Day 14 vs. day 28 | 0.02269 | -0.4228 to 0.4682 | ns | >.999 | 15 | 15 |
| Day 28 vs. day 42 | 0.009367 | -0.5483 to 0.5670 | ns | >.999 | 15 | 15 |
| veh + CGRP 0.01 mg kg-1 | | | | | | |
| B vs. day 1 | 0.9332 | 0.5160 to 1.350 | *** | <.001 | 29 | 29 |
| Day 1 vs. day 14 | 0.09078 | -0.2462 to 0.4277 | ns | 0.914 | 15 | 15 |
| Day 14 vs. day 28 | 0.01803 | -0.02702 to 0.06309 | ns | 0.725 | 15 | 15 |
| Day 28 vs. day 42 | -0.00036 | -0.004817 to 0.004097 | ns | 0.999 | 15 | 15 |
| PZN + veh | | | | | | |
| B vs. day 1 | 0.8233 | 0.1998 to 1.447 | ** | 0.009 | 12 | 12 |
| Day 1 vs. day 14 | -1.148 | -1.663 to -0.6322 | *** | <.001 | 12 | 12 |
| Day 14 vs. day 28 | 0.06164 | -0.7093 to 0.8325 | ns | 0.999 | 12 | 12 |
| Day 28 vs. day 42 | -0.04877 | -0.7797 to 0.6822 | ns | >.999 | 12 | 12 |
| PZN + CGRP 0.01 mg kg-1 | | | | | | |
| B vs. day 1 | 0.8233 | 0.1998 to 1.447 | ** | 0.009 | 12 | 12 |
| Day 1 vs. day 14 | -0.004972 | -0.04848 to 0.03854 | ns | 0.995 | 12 | 12 |
| Day 14 vs. day 28 | 0.02058 | -0.07620 to 0.1174 | ns | 0.955 | 12 | 12 |
| Day 28 vs. day 42 | 0.007808 | -0.005733 to 0.02135 | ns | 0.388 | 12 | 12 |

### Figure 8I

Kruskal-Wallis test

| | |
|---|---|
| Number of families | 1 |
| Number of comparisons per family | 3 |
| Alpha | 0.05 |

| | |
|---|---|
| P value | <.001 |
| Exact or approximate P value? | Approximate |
| P value summary | *** |
| Do the medians vary signif. (P < 0.05)? | Yes |
| Number of groups | 4 |
| Kruskal-Wallis statistic | 42.78 |

| Dunn's multiple comparisons test | Mean rank diff. | Significant? | Summary | Adjusted P Value | n1 | n2 |
|---|---|---|---|---|---|---|
| veh + veh vs. veh + CGRP 0.01 mg kg-1 | 25.33 | Yes | *** | <.001 | 15 | 15 |
| PZN + veh vs. PZN + CGRP 0.01 mg kg-1 | 29.08 | Yes | *** | <.001 | 12 | 12 |
| veh + CGRP 0.01 mg kg-1 vs. PZN + CGRP 0.01 mg kg-1 | 0.825 | No | ns | >.999 | 15 | 12 |

### Figure 8J

Kruskal-Wallis test

| | |
|---|---|
| Number of families | 1 |
| Number of comparisons per family | 3 |
| Alpha | 0.05 |

| | |
|---|---|
| P value | <.001 |
| Exact or approximate P value? | Approximate |
| P value summary | *** |
| Do the medians vary signif. (P < 0.05)? | Yes |
| Number of groups | 4 |
| Kruskal-Wallis statistic | 45.81 |

| Dunn's multiple comparisons test | Mean rank diff. | Significant? | Summary | Adjusted P Value | n1 | n2 |
|---|---|---|---|---|---|---|
| veh + veh vs. veh + SNP 0.25 mg kg-1 | 27.13 | Yes | *** | <.001 | 15 | 15 |
| PZN + veh vs. PZN + SNP 0.25 mg kg-1 | 26.67 | Yes | *** | <.001 | 12 | 12 |
| veh + SNP 0.25 mg kg-1 vs. PZN + SNP 0.25 mg kg-1 | -4.283 | No | ns | >.999 | 15 | 12 |
